# On the Distribution of Free-Energy in Metabolism

**DOI:** 10.64898/2026.06.01.729314

**Authors:** Ian Cook, Thomas S. Leyh

**Author notes:** Corresponding Author Address:* The Department Microbiology and Immunology, Albert Einstein College of Medicine, 1300 Morris Park Ave., Bronx, New York 10461-1926.

## Abstract

Chemical potential is coupled to cellular processes by the flow of metabolites through catalytic networks known collectively as metabolism. Here we describe an extensive new class of energy-coupling catalysts that act to interconnect metabolic network pathways and their potentials. Members of the class are defined by a common mechanism — half-site reactivity. The well-established sequential subunit turnover of half-site enzymes suggests that the potentials of reactions occurring at the separate subunits are coupled to one another. Here this hypothesis is tested and validated using promiscuous half-site enzymes from two catalytically distinct enzyme families, each with broad metabolic penetrance. Fundamental catalytic parameters (V_max_ and K_m_) and reaction endpoints are predicted and shown to change dramatically when reaction potentials are coupled — for example, the catalytic efficiency (V_max_/K_m_) and endpoint of the retinol oxidation reaction (the rate-limiting step in vitamin A synthesis) are shown to increase 900- and 3,400-fold, respectively, when the reaction is coupled to the more favorable oxidation of ethanol. For the first time it is clear that metabolism has the flexibility to react to changes in the metabolic state of the cell by redistributing chemical potential among the many metabolic pathways interconnected by half-site enzymes.

**Significance:** The findings herein reveal the existence of an extensive, catalytically diverse network of enzymes that distributes chemical potential within and across the pathways of small-molecule metabolism. Members of the network are identified on the basis of a shared mechanistic trait — half-site reactivity. These energy-coupling catalysts allow reactions to proceed orders of magnitude further and more efficiently than their intrinsic potentials allow by coupling them to more favorable reactions. The work offers a *raison d’etre* for the half-site mechanism and powerful new strategies for *de novo* metabolic pathway design.

---

Chemical potential is arguably the most precious resource of the cell — without it, all molecular processes would “stand still,” and life as we know it would cease. The molecular machines that mediate the accumulation and distribution of chemical potential in metabolism continue to be a subject of intense interest^1^. Work in the 1950s advanced the concept of metabolic vectorial processes by demonstrating that an osmotic gradient — in effect, a cellular battery — could be established by coupling ATP hydrolysis to ion-transport across a membrane^2^. In the 1970s it was recognized that the potentials of such seemingly distinct processes can be linked to one another *via* conformational changes in the proteins that couple them^3^. These energy-coupling changes are elicited by catalytic-stage dependent allosteric interactions that stoichiometrically link the processes. In recent decades, the energetics of a stunning array of conformationally coupled biological processes have been discovered^4-10^ and elucidation of their molecular mechanisms has led to the development of man-made nanomachines^11,12^. Here we demonstrate that half-site enzymes — a mechanistic class of enzymes found throughout metabolism — are energy-coupling catalysts that link the chemical potentials of the reactions they catalyzed.

As the namesake implies, only half of the subunits of a half-site enzyme appear to be reactive. Numerous studies offer intriguing discussions regarding the mechanistic basis of half-site reactivity^13-15^; however, to our knowledge, a *raison d’etre* for such systems has not been proposed. Our findings suggest that by intertwining catalytically competent, sequence-identical monomers such that the subunits of the resulting oligomers turnover in a step-wise fashion, Nature has provided the selective advantages associated with harnessing metabolic-reaction potentials to one another.

Half-site reactivity is often taken as evidence of sequential subunit turnover. Such a mechanism requires inter-subunit allosteric communications that stoichiometrically link the subunit catalytic cycles, which implies that the potentials of the individual subunit-catalyzed reactions are coupled. Here we demonstrate that half-site enzymes do indeed couple the potentials of the reactions they catalyze. Further, subunit catalytic efficiency is shown to be a sensitive function of the difference in the potential of the coupled reactions; hence, initial-rate parameters can be manipulated. The metabolic prevalence of half-site enzymes implies the existence of an extensive catalytic network that distributes chemical potential across small-molecule metabolic pathways and imbues metabolism with thermodynamic and kinetic flexibility it would not otherwise have.

## Results and Discussion

### The Theory

Assuming lossless coupling, the chemical potential shared by two coupled reactions — the *interaction energy* (ΔG_*int*_) — is given by the difference in the potentials of the individual reactions; hence,

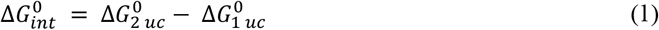

where the *uc* subscript indicates an uncoupled reaction, and the naught superscript indicates a standard free energy.

Upon coupling, the potential of the less favorable reaction increases by the interaction energy, and, by conservation-of–energy, that of the more favorable reaction decreases by an equivalent amount; thus,

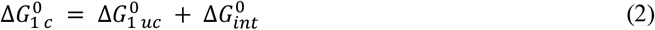

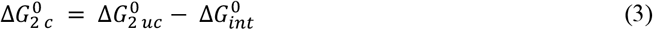

where the subscript *c* indicates a coupled potential. Applying the Nernst equation,

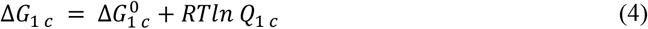

and substituting equation 2,

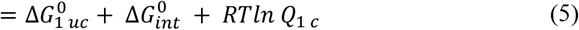

where *Q*_*1*_ is the mass-action ratio (product/substrate) of Reaction 1, and *R* and *T* are the ideal gas constant and absolute temperature.

At equilibrium,

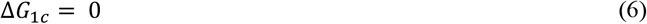

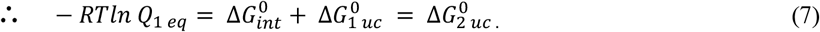

Equation 7 predicts that the mass-action ratio of the coupled Reaction 1 at equilibrium (*Q*_1*c*_) is given by the uncoupled potential of the reaction to which it is coupled (i.e., 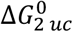). Thus, the equilibrium distribution of either coupled reaction can be predetermined by judicious selection of the partnered reaction potential.

The ramifications of Equation 7 can be couched in terms of enzyme initial-rate parameters. The catalytic efficiency of an enzyme (k_cat_/K_m_) is related to reaction potential by the Haldane relationship^16^, which expresses the reaction equilibrium constant in terms of the enzyme’s initial-rate parameters. The Haldane governing interconversion of a single substrate (A) and product (P) is given by Equation 8:

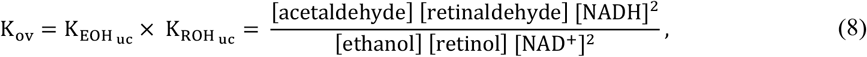

where *f* and *r* indicate forward and reverse reactions. As a reaction shifts from an uncoupled to a coupled condition, its equilibrium reactant distribution shifts in accordance with Equation 7, and its catalytic efficiency must also shift to maintain the Haldane relationship. Hence, catalytic efficiencies, like equilibrium endpoints, are manipulable — the efficiency toward a given substrate is set by the potential of the reaction to which it is coupled.

The Haldane for a two-reaction system (*A-to-P* and *B-to-Q*) coupled in 1:1 stoichiometry can be written:

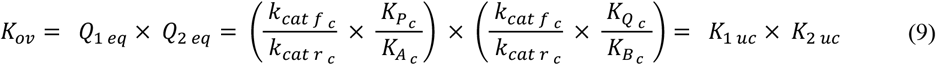

where the system’s overall equilibrium constant, *K*_*ov*_, is given by the product of the reaction equilibrium mass-action ratios, *Q*_*eq*_, and by the product of the uncoupled equilibrium constants, *K*_*uc*_. The functions in the right- and left-hand parentheses correspond to *Q*_1 *eq*_ and *Q*_1 *eq*_, respectively. Notably, for stoichiometrically coupled (1:1) reactions the forward k_cat_ values are identical, as are the reverse k_cat_ values.

### Testing the Theory

Central to the coupling theory is the prediction that the mass-action ratio of a perfectly coupled reaction (i.e., lossless transfer of potential) at equilibrium is given by the Gibbs potential of the reaction to which it is coupled (see, Equation 7). A corollary prediction is that the catalytic efficiencies of coupled reactions will adjust to satisfy their equilibrium mass-action ratios. These predictions are validated below in a series of initial-rate and equilibrium measurements using representative half-site enzymes from two enzyme families —alcohol dehydrogenases (ADHs) and glutathione S-transferases (GSTs).

Half-site enzymes can form *homocomplexes*, which couple identical reactions to one another, and *heterocomplexes*, which couple different reactions. When the potentials of the reactions they catalyze differ, heterocomplexes catalyze a net transfer of potential. Homocomplexes couple reactions with identical reactions potentials; hence, they produce zero net transfer of potential and provide a path for the potential transferred by heterocomplexes to “leak” from the system. Minimizing the homocomplex/ heterocomplex ratio slows the leak and increases coupling efficiency. The specificities of heterodimer subunits often differ considerably and judicious matching of substrates and subunit specificities can be used to maximize heterocomplex formation. ADHs and GSTs form hetero- and homodimers *in vivo* and both are used in our studies^17,18^.

### ADH Studies

ADH1s are dimeric, found in gram quantities in adult liver, and form heterodimers composed of any two of four possible subunit types (α, β, and γ). The ADH1 α- and γ-isozymes^19^ are promiscuous^20^, half-site reactive^21,22^ and can be low-temperature induced to form monomers^23^ from which stable heterodimers can be constructed (*Materials and Methods*). Among ADH1 isoforms, ADH1α has the highest catalytic efficiency towards retinol^24^ and lowest towards ethanol, and ADH1γ efficiencies are the reverse^24^.

Initial-rate parameters for the individual retinol and ethanol reactions were determined in the forward and reverse directions for αα−, γγ− and αγ-ADH1, and parameters for the coupled-reaction were determined for the αγ form. Homo- and heterodimer parameters are compiled in Tables 1 and 2, respectively. Initial-rate protocols are outlined in *Material and Methods*, and experimental conditions are given in the Legends of Figs S1-S4, which present the initial-rate data and fitting results.

**Table 1.** Initial-Rate Parameters and Equilibrium Constants for ADH1 Homodimer Catalyzed Reactions.

|  | Ethanol Reaction |  |  |  | Retinol Reaction |  |  |  |
| --- | --- | --- | --- | --- | --- | --- | --- | --- |
|  | *Forward* |  | *Reverse* |  | *Forward* |  | *Reverse* |  |
|  | *αα*-Homodimer |  |  |  | *αα*-Homodimer |  |  |  |
| Substrate | NAD<sup>+</sup> | EOH | NADH | Acetaldehyde | NAD<sup>+</sup> | Retinol | NADH | Retinal |
| K<sub>m</sub> (μM) | 210 (12)<sup>a</sup> | 3700 (210) | 68 (4.1) | 260 (16) | 310 (25) | 45 (2.1) | 1.1 (0.07) | 0.46 (0.038) |
| k<sub>cat</sub> (min<sup>-1</sup>) | 120 (7) |  | 4300 (280) |  | 62 (3.1) |  | 3100 (250) |  |
|  | *γγ*-Homodimer |  |  |  | *γγ*-Homodimer |  |  |  |
| K<sub>m</sub> (μM) | 280 (15) | 120 (7) | 13 (1.6) | 12 (1.5) | 230 (19) | 190 (15) | 2.5 (0.11) | 0.47 (0.025) |
| k<sub>cat</sub> (min<sup>-1</sup>) | 150 (11) |  | 1100 (140) |  | 45 (3.6) |  | 1700 (120) |  |
| K<sub>eq</sub> Haldane<sup>b</sup> | 6.3 x 10<sup>-4</sup> (1.1 x 10<sup>-4</sup>) |  |  |  | 7.2 x 10<sup>-7</sup> (1.4 x 10<sup>-7</sup>) |  |  |  |
| K<sub>eq</sub> exp<sup>c</sup> | 7.0 x 10<sup>-4</sup> (0.5 x 10<sup>-5</sup>) |  |  |  | 6.8 x 10<sup>-7</sup> (0.4 x 10<sup>-8</sup>) |  |  |  |
| K<sub>ov</sub><sup>d</sup> | 4.5 x 10<sup>-10</sup> (1.1 x 10<sup>-10</sup>) |  |  |  |  |  |  |  |

<sup>a</sup>Values in parentheses indicate one standard deviation unit. <sup>b</sup>K<sub>eq</sub> Haldane, value calculated from initial-rate parameters using the Haldane Relationship. <sup>c</sup>K<sub>eq</sub> exp, value determined from measurements at equilibrium. <sup>d</sup>K<sub>ov</sub>, the product of the EOH- and ROH-reaction K<sub>eq</sub> Haldane values — the predicted coupled-reaction equilibrium constant.

**Table 2.**
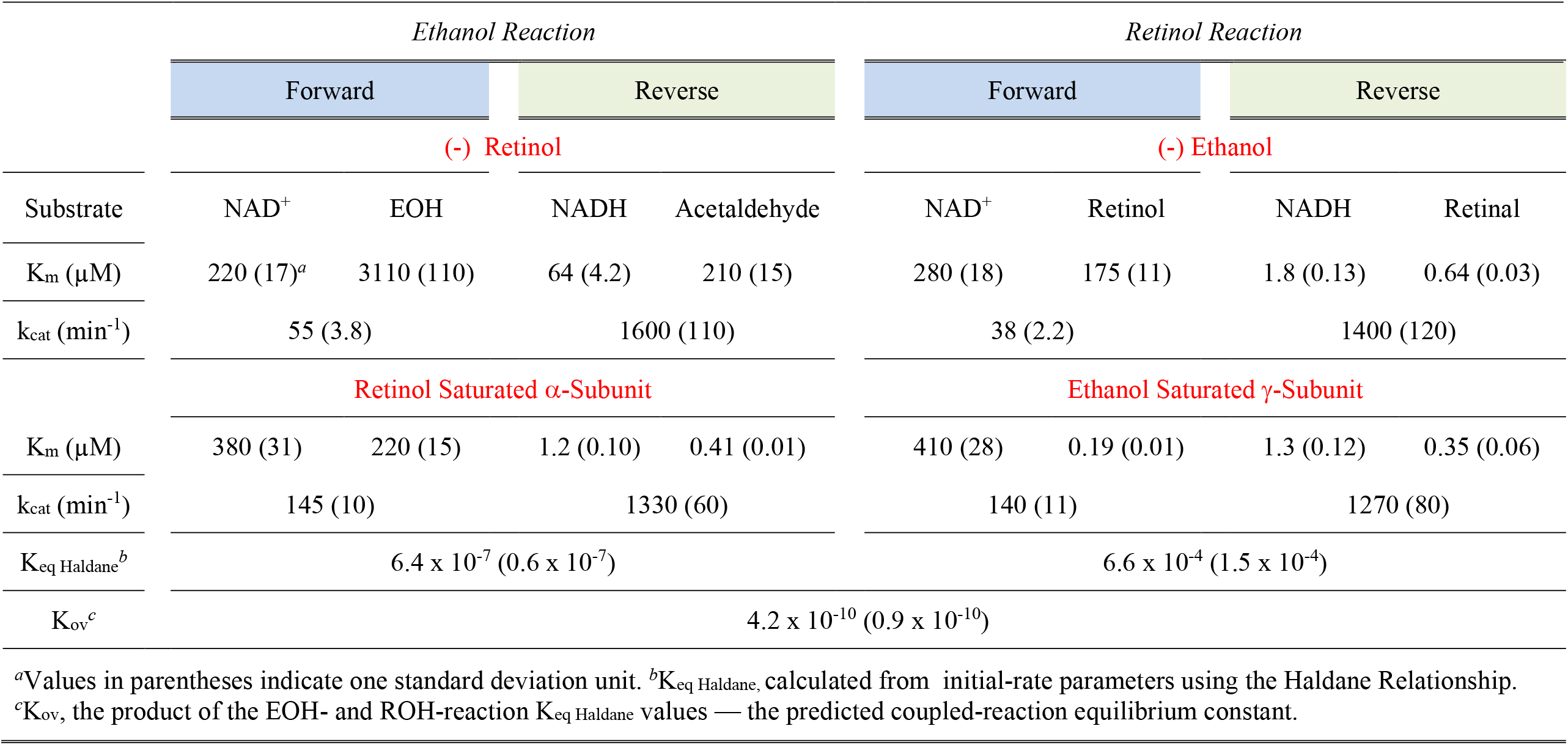
Initial-Rate Parameters and Equilibrium Constants for αγ-ADH1 Heterodimer Catalyzed Reactions.

The Table 1 parameters reveal a clear preference of the γγ-over the αα-isoform for the EOH reaction and a lesser preference of the αα-over the γγ-isoform for the ROH reaction. The Table’s bottom rows contain equilibrium constants for the individual reactions, calculated both from measurements at equilibrium (K_eq exp_) and the Haldane Equation (K_eq Haldane_), and the “overall” equilibrium constant, K_ov_ — the product of K_eq Haldane_ for both reactions. It is notable that the heterodimer parameters for the individual reactions (Table 2) resemble those of the less efficient homodimer (Table 1) — a finding consistent with a sequential-turnover mechanism in which the catalytic cycle cannot be completed until the low-affinity subunit has turned over.

The coupled-reaction initial-rate parameters were obtained under conditions where ≥ 98% of the enzyme is in heterocomplex and thus report the behavior of the system operating at ~ 100% efficiency. Remarkably, the coupled ethanol reaction parameters have shifted to closely mimic those of the uncoupled retinol reaction (Tables 1 and 2). This shift is also apparent in the equivalence of the equilibrium constants calculated for the coupled ethanol (Table 2) and uncoupled retinol (Table 1) reactions 6.4 ± 0.6 × 10^−7^ and 7.2 ± 1.4 × 10^−7^. Similarly, the efficiencies of the coupled retinol reaction (Table 2) have shifted such that its energetics match those of the uncoupled ethanol (Table 1) reaction (6.6 ± 1.5 × 10^−4^ and 6.3 ± 1.1 × 10^−4^). As predicted by the theory, the catalytic efficiencies have shifted such that the Gibbs potential of a given reaction, calculated from the Haldane, has become equal to that of its partnered reaction. The reaction energetics have counterbalanced such that the total energy in the system is maintained.

To further confirm the half-site coupling hypothesis, coupling was evaluated at equilibrium. The equilibrium studies involved allowing the uncoupled ethanol reaction to equilibrate prior to the addition of retinol and a subsequent second equilibration. The uncoupled ethanol reaction equilibrium constant calculated from the first equilibration, 6.5 ± 0.4 × 10^−4^, is equal to that calculated using the Haldane, 6.3 ± 1.1 × 10^−4^, Table 1. The addition of retinol, at 1 hr, causes a transient burst of products (see, Fig1B) after which the reactions re-equilibrate.

Unlike the initial-rate studies, measurements at equilibrium are taken when the system is operating at less than 100% efficiency due to increased contributions from homocomplex pathways, which do not catalyze a net transfer of potential. As homocomplex levels increase, the potential accumulated in the earlier stages of reaction, where efficiency is higher, “leaks” from the coupled reactions, causing them to shift toward their uncoupled equilibrium positions. The total energy of the system is conserved regardless of transfer efficiency; hence, at equilibrium:

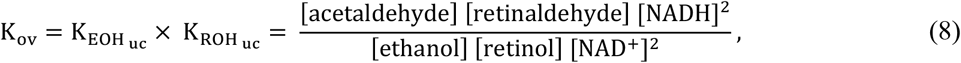

where K_EOH uc_ and K_ROH uc_ are the uncoupled-reaction equilibrium constants. The K_ov_ values calculated using either the uncoupled-reaction equilibrium constants (Table 1) or the reactant concentrations in the plateau of the second-phase reaction (Fig 1) are identical within error (4.5 × 10^−10^ and 4.4 × 10^−10^, respectively). The potential transferred between the reactions can be calculated by subtracting from the Gibbs potential of either reaction (calculated using reactant levels in the second plateau) the uncoupled-reaction potential of that same reaction (Eq 1). Such calculations yield a net transfer of 2.0 kcal/mole to the ethanol reaction, and −2.0 kcal/mole to the retinol reaction. Consistent with conservation-of-energy, transfer is equal and opposite — the potential “gained” by one reaction is “lost” by the other. The maximum potential that can be transferred (which occurs at 100% transfer efficiency) is calculated at 4.1 kcal/mole (see, Eq 1); hence, the transfer efficiency under these conditions is 49%.

**Figure 1.**
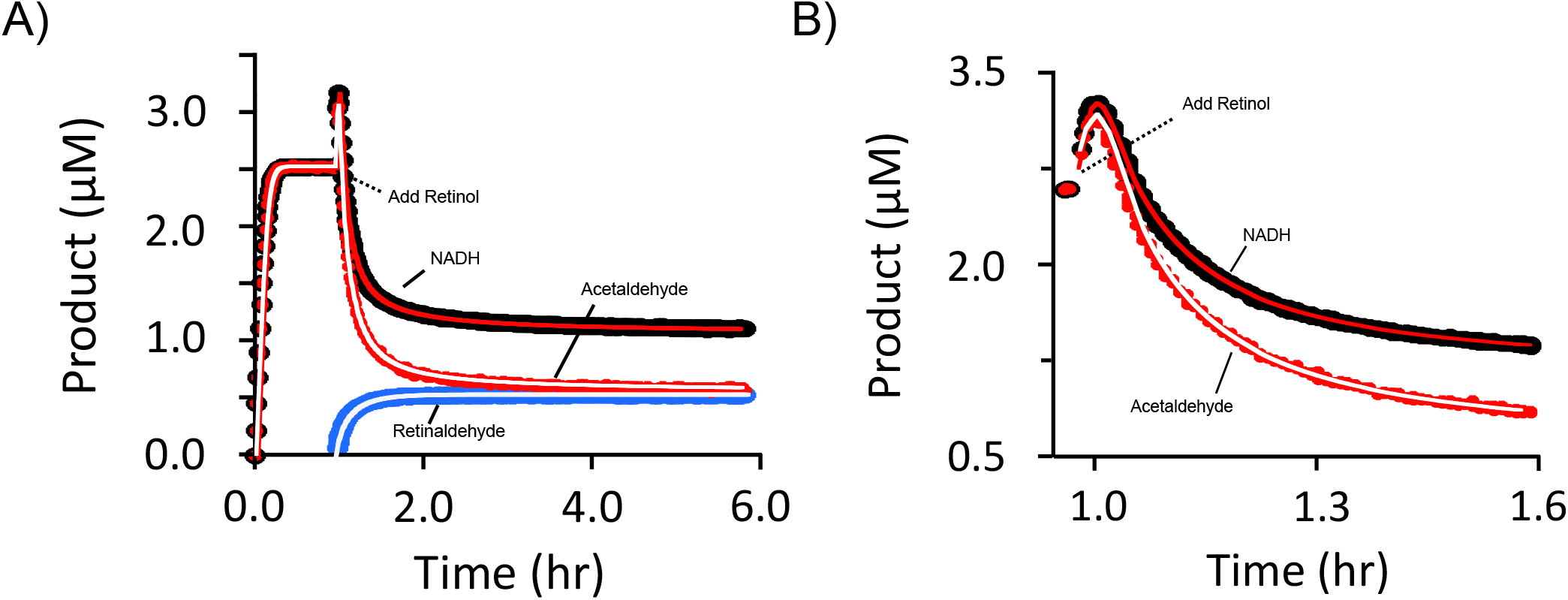
ADH Heterodimer Coupling at Equilibrium. (A) *The ROH/EOH-Coupled Reactions*. The α,γ-ADH1 catalyzed uncoupled ethanol-oxidation reaction was initiated by addition of NAD^+^ and allowed to reach equilibrium. The coupled reaction was then initiated by adding retinol (at 1.0 h) and allowed to equilibrate. Acetaldehyde, retinaldehyde and NADH levels are represented by red, blue, and black lines, respectively. Retinaldehyde and NADH concentrations were monitored continuously *via* fluorescence (λ_ex_ = 420 nm, λ_em_ = 530 nm) and absorbance (ε_340_ = 6.22 mM^−1^ cm^−1^), respectively; acetaldehyde is given by [NADH] – [RA]. Reaction conditions at t_0_: α,γ-ADH1 (0.10 nM, heterodimer), NAD^+^ (10 mM, 25 x K_m_), ethanol (4.0 µM), KPO_4_ (50 mM), pH 7.4, 25 ± 2ºC. At one hour, retinol was added to 4.0 µM. Reactions were performed in triplicate; the averaged data are shown. Lines passing through the data represent the behavior predicted by the coupled-ADH model parameterized using the initial-rate constants in Tables 1 and 2 (see, *Supplement 1, Modeling*). (B) *The Pulse Following Retinol Addition (Expanded Panel A Time Axis)*.

**Figure 2.**
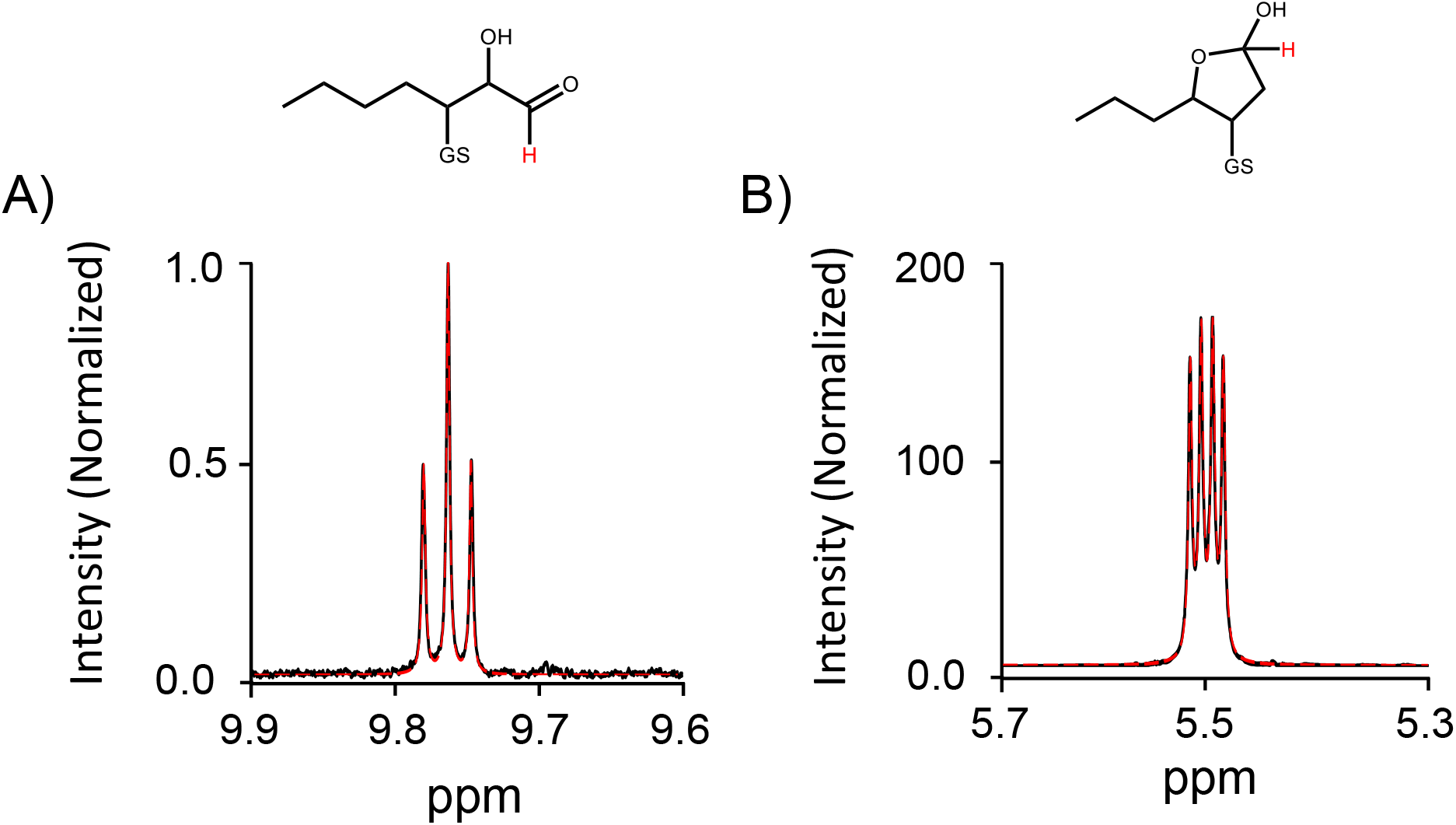
Linear and Cyclical GS-4HNE H1-Proton NMR Spectra. Structures and H1-proton spectra of the GS-4HNE linear and cyclical forms are given in Panels A and B. H1 protons are indicated by red text. The maximum peak height in each spectrum is normalized to the Panel A peak-height maximum. Dashed red lines indicate the fits used to obtain the peak areas and isomerization equilibrium constant. NMR parameters and solution conditions are given in *Methods, GS-4HNE* ^*1*^*H-NMR*.

The impact of coupling on α,γ-ADH1 catalytic efficiency and reaction energetics is remarkable, but no more so than the simple fact that these parameters, long considered immutable, can be predictably manipulated by suitable selection of reaction partners. Consider the retinol reaction… its energetics become ~1000-fold more favorable upon coupling, K_m retinol_ decreases ~900-fold to 200 nM (a value comparable to unbound retinol concentration in human serum^25^) and the catalytic efficiency (k_cat_/K_m_) of the enzyme, which, along with other promiscuous ADH isoforms, catalyzes the rate-limiting step in vitamin A biosynthesis^26^, increases 3400-fold toward retinol.

More than 140,000 unique ADH sequences are archived in public databases^27^ and the associated enzymes engage in myriad metabolic processes including, in humans, vision^28^, the production and elimination of toxins^29,30^, and the synthesis and regulation of hydroxysteroids^31^, prostaglandins^32^ and neurotransmitters^33^. The ADH superfamily is structurally conserved and composed of numerous promiscuous dimers belonging to subfamilies that contain half-site reactive members. In total, these facts suggest that the coupling of ADH reactant potentials occurs throughout the biome.

### Half-Site Coupling in the Glutathione S-Transferase Family

To demonstrate that energy coupling is a property of half-site mechanisms rather than a unique feature of the ADH family, parallel initial-rate and equilibrium studies were performed with half-site enzymes from a second family — the glutathione S-transferases (GSTs).

GSTs play important roles in neutralizing toxins^34,35^, metabolite synthesis^36^, and as reactive-oxygen and -nitrogen chemosensors^34^. Toxins are neutralized by GST-catalyzed nucleophilic attack of the glutathione thiolate anion at their electrophilic centers to form non-toxic GST conjugates^37,34,35^. GST substrate specificities are typically remarkably broad^38^ and center on distinct metabolic areas. Humans express 16 isoforms, which are organized into seven classes. The alpha class contains five isoforms GSTA1-GSTA5, which are expressed primarily in liver and found both as hetero- and homodimers *in vivo* (). Our studies pair the human GSTA1 and GSTA4 isoforms with the substrates 1,2-dichloro-4-nitrobenzene (DCNB), a well-known environmental toxin^39^, and 4-hydroxy-nonenal (4HNE), a disease-linked by-product of lipid peroxidation^40^.

Proper interpretation of the initial-rate and equilibrium findings described below requires careful consideration of the fact that the GS-4HNE conjugate undergoes a thermodynamically favorable non-enzymatic cyclization reaction ^41^. The linear and cyclic forms of GS-4HNE are distinguishable based on ^1^H-NMR of the H1 proton, which undergoes a significant change in chemical shift (−4.48 ppm) as GS-4HNE moves from the linear (aldehyde) to cyclic (hemiketal) form. Both H1 peaks are baseline isolated from other resonances (see, Fig X). K_eq_ calculated from H1 peak areas obtained using TopSpin^42^ is 180 ± X in favor of the cyclized form. The value of 180 was independently corroborated by taking the ratio of the forward and reverse cyclization rate constants determined using H1 saturation-transfer (see, *Supplement 1, GS-4HNE* ^*1*^*H-NMR*). The forward and reverse rate constants, 3100 s^−1^ and 17 s^−1^, predict a K_eq_ of 182 ± 24. Given that the linear form of 4HNE is the enzyme substrate (PDB: 3IK7^43^), the experimentally determined K_m GS-4HNE_ and K_eq_ values were decreased by a factor of 180 (see, Table 3 and 4 footnotes).

**Table 3.**
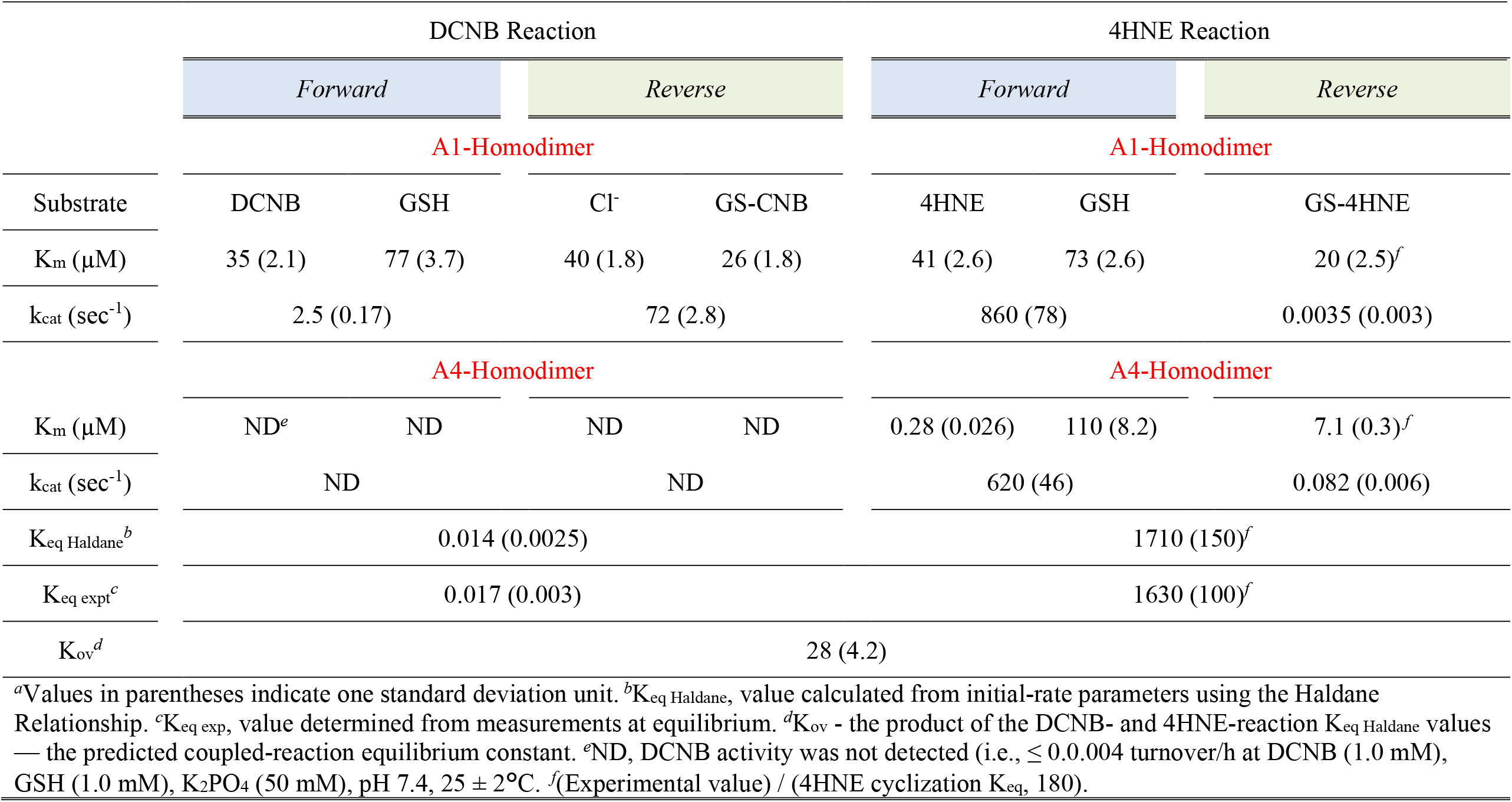
Initial-Rate Parameters and Equilibrium Constants for GST Homodimer Catalyzed Reactions.

**Table 4.**
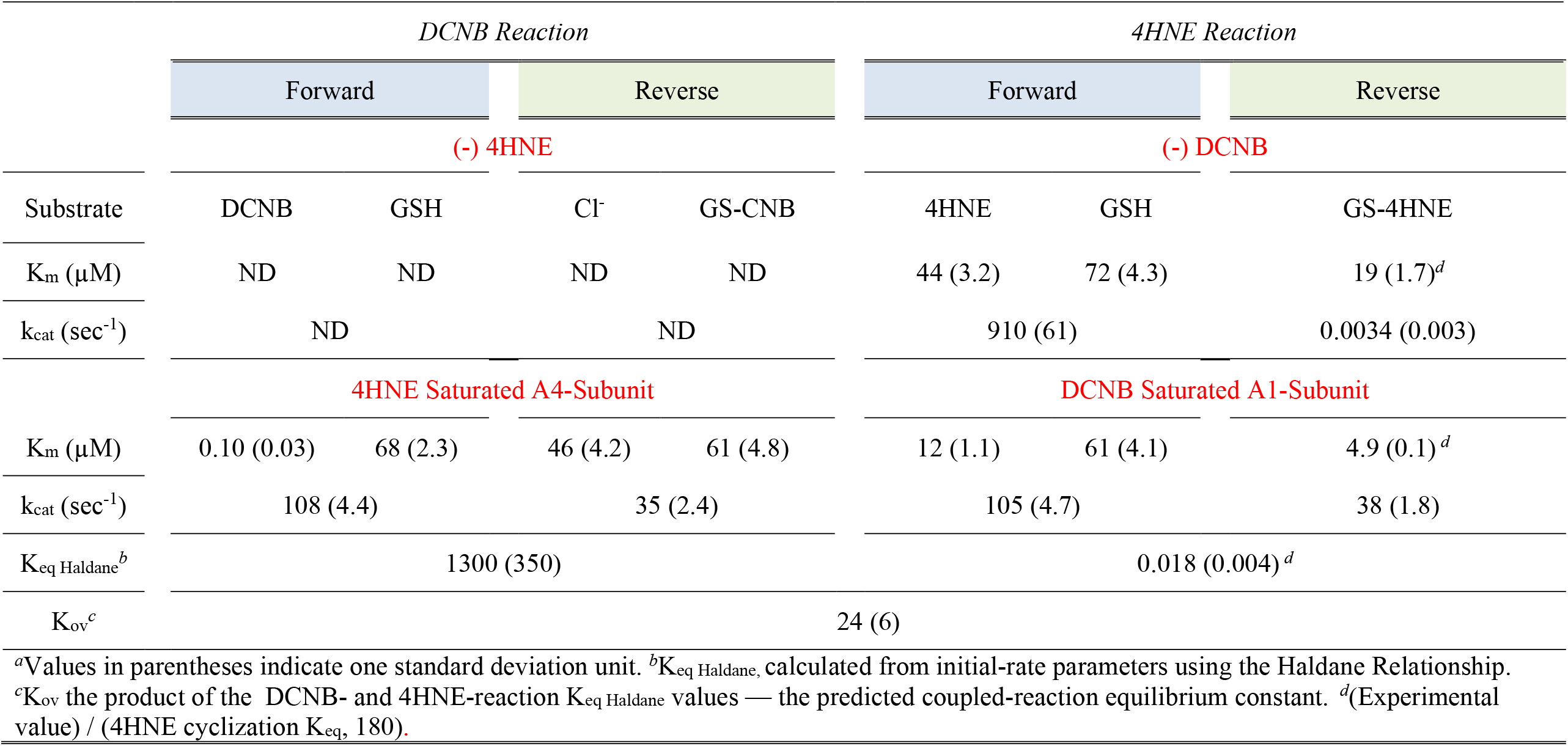
Initial-Rate Parameters and Equilibrium Constants for the GSTA1/A4 Heterodimer Catalyzed Reactions.

Initial-rate parameters for the forward and reverse DCNB- and 4HNE-conjugation reactions were determined for the GST homo- and heterodimers. The parameters, compiled in Tables 3 and 4, agree well with literature values^43-47^. The associated experimental protocols and primary data are provided in *Supplementary Materials* and Figs S5-8. In the absence of DCNB, the GSTA1 homo- and heterodimer 4HNE-reaction parameters are indistinguishable, while those of the GSTA4 homodimer differ markedly. That the 4HNE parameters of the heterodimer and GSTA1 homodimer are indiscernible suggests that the A1-subunit behavior in the heterodimer is little influenced by that of the A4 subunit, and vice versa. However, as expected, the subunit parameters shift once the heterodimer is coupling reactions with different potentials.

Equilibrium constants for the individual DCNB and 4HNE reactions were determined by allowing the reactions to run to completion. To ensure reaction endpoints were not due to enzyme inactivation or cryptic inhibition, substrate was added following the first endpoint plateau, and a second plateau was then reached. Each dual-plateau experiment yields two K_eq expt_ measurements, which agreed in all cases. Three such experiments were performed for each reaction, yielding six measurements that were averaged obtain K_eq expt_. The resulting DCNB and 4HNE reaction equilibrium constants (0.014 ± 0.0025 and 1710 ± 150, respectively) agree well with the constants calculated from the initial-rate parameters using the Haldane equation (0.017 ± 0.003 and 1630 ± 100), see Table 3. The Cl^−^ concentration in the reaction buffer was determined and taken into account in the equilibrium constant calculations. Experimental protocols and data associated with the K_eq expt_ studies are given in *Supplemental Materials, Equilibrium Constant Determinations*, and Fig S9.

To assess coupling of the DCNB and 4HNE reactions at equilibrium, a comparative study was performed in which reactions were allowed to reach equilibrium under identical conditions except that they were either “coupled,” using the GSTA1/A4 heterodimer (300 μM), or “uncoupled,” using the GSTA1 and A4 homodimers (150 μM, each). Results and conditions are given in Fig 3 and its legend. Each datapoint is the average of three determinations and lines passing through the data represent the behavior predicted by the GST coupling model (see, *Supplement 1, Modeling*) using the initial-rate constants in Tables 3 and 4. As is apparent (Fig 3A and B) DCNB conjugation is driven substantially forward by the presence of the heterodimer. K_eq_ for the homo- and heterodimer catalyzed reactions shifts from 0.012 ± 0.004 to 380 ± 80 — a remarkable 32,000-fold increase. Conversely, 4HNE conjugation undergoes a 30,000-fold decrease in its equilibrium position as a consequence of coupling — 1800 ± 150 to 0.059 ± 0.012. The efficiency of free energy transfer, taken as the ratio of the experimentally observed shift (30,000-fold) to that calculated at 100% efficiency (140,000-fold), is 21%.

**Figure 3.**
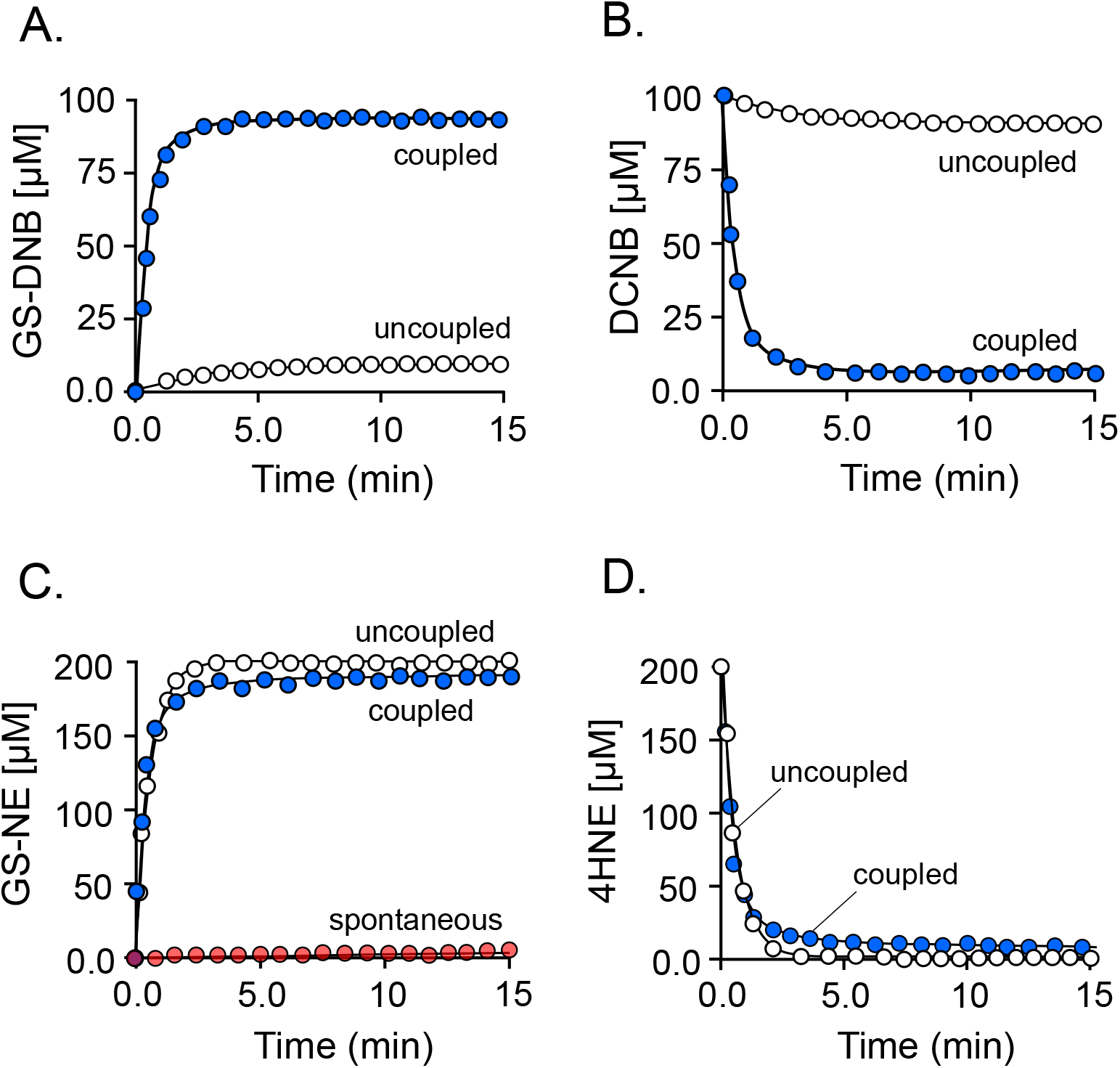
Coupled and Uncoupled GST-Conjugation Reactions. **(A and B)** *Coupled and uncoupled, forward* (**A**) *and reverse* (**B**) *DCNB conjugation reactions*. **(C and D)** *Coupled and uncoupled, forward* (**C**) *and reverse* (**D**) *4HNE conjugation reactions*. Panel A-D reaction conditions: DCNB (300 µM, 59 x K_m_), 4HNE (200 µM, 4.4 x K_m_), GSH (300 µM, 4.0 x K_m_), DTT (2.0 mM), KPO_4_ (50 mM), pH 7.4, 25 (±2) °C. Coupled reactions contained GSTA1/A4 heterodimer (300 nM, dimer). Uncoupled reactions contained a 1:1 mixture of homodimers, GSTA1 (150 nM) and GSTA4 (150 nM). Forward and reverse Reactions were monitored *via* the absorbance of DCNB (ε_410_ = 9600 M^−1^ cm^−1^), GS-DNB (ε_345_ = 8500 M^−1^ cm^−1^), 4HNE (ε_225_ = 19200 M^−1^ cm^−1^) and GS-NE (ε_285_ = 22000 M^−1^ cm^−1^). Reactions were initiated by addition of GSH. Each progress curve is the average of three reactions. Lines through the points show the behavior predicted using kinetic constants in Tables 3 and 4. Progress of the 4HNE conjugation reaction was monitored in the absence of enzyme (red dots) to ensure that the spontaneous conjugation reaction did not significantly contribute to the measurements.

These finding corroborate the ADH studies and independently confirm that half-site systems couple the potentials of the reactions they catalyze. Moreover, here we see two human toxins profoundly influencing one another’s reactivities and reaction endpoints — such interplay is expected to extend to drug-drug interactions.

### Implications

The human ADH1 family consists of four distinct isoforms capable of producing ten different dimers, eight of which have thus far been purified from liver^18^. Given the many metabolic pathways in which ADH1 dimers operate, scores of hepatocyte metabolites will likely compete for the ADH active sites. Such environments favor the formation of heterocomplexes, which will accelerate and drive unfavorable reactions further than their intrinsic potentials allow at the “expense” of more favorable reactions. In such situations, the driving force of a given reaction is a complex function of the aggregate potential of the reactions to which it is coupled. Promiscuous half-site enzymes link the potentials of all the pathways in which they participate, which raises the question of whether such linked pathways are better viewed as linear concatenates of stepwise reactions with discrete potentials, or as multidimensional networks interconnected at the steps coupled by half-site enzymes.

From a pragmatic perspective, perhaps the most valuable finding of this study is that the initial-rate parameters and extent of a reaction can be modulated by judicious choice of the partnered reaction. Flexibility in these variables provides a powerful means of controlling both the flux and yields of metabolic pathways. Pathway design lies at the heart of the bioengineering industry, which is continually developing pathways to enrich foodstuffs^48,49^ and produce fertilizers^50^, biofuels^51^, natural-products^52^, drugs^53,54^, and fine chemicals^55^.

Molecular motors couple small-molecule chemistry to vectorial macromolecular motions that lead to mechanical outcomes. Half-site systems, which couple small-molecule chemistries to one another, also depend on vectorial motion — they oscillate in fixed sequence through a distinct series of conformational states at a frequency linked to the turnover of the enzyme. It is plausible that half-site enzymes can be evolved into nanomachines by adapting their oscillating vectorial motions to accomplish mechanical tasks.

The majority of enzymes have not been tested for half-site reactivity. Typically, such systems are discovered serendipitously when, under a particular condition, one observes that only one half of the subunits appear to be reactive. The current literature reports twenty-seven half-site containing enzyme families (*Supplement 2*) each with its own unique structural characteristics, distinct chemistry and metabolic niche. Over evolutionary time, Nature has entwined what are often catalytically competent monomers in these families such that their mechanisms are interdigitated and the potentials of the chemistries they catalyze are coupled. It seems much can be learned from studying the ways in which Nature has solved the structural/mechanistic puzzle of transforming identical monomers into to energy-coupling dimers.

While we have yet to fully understand the advantages that half-site enzymes bring to systems as complex as human metabolism, which is comprised of ~8000 reactions^56^, we now know that half-site reactivity imbues metabolism with a kinetic and energetic flexibility that enables it to both shape and respond to the metabolic poise of the cell. From an evolutionary vantage point, oligomerization events that lead to half-site mechanisms enable metabolic systems to incorporate reactions that are otherwise too inefficient and unfavorable to be metabolically viable.

### Conclusions

1, Half-site enzymes couple the chemical potentials of the reactions they catalyze; and 2, The endpoints and initial-rate parameters of coupled reactions are selectable and determined by the potentials of their reaction partners.

## Materials and Methods

### Materials

Acetaldehyde, acetic acid, ammonium hydroxide, 1-Chloro-2,4-dinitrobenzene (CDNB), dibasic potassium phosphate, 1,2-dichloro-4-nitrobenzene (DCNB), 5,5-dithiobis (2-nitrobenzoic acid) (DTNB), dithiothreitol (DTT), ethanol (200 proof), ethylenediaminetetraacetic acid (EDTA), L-glutathione (reduced), (2-cholro-4-dinitrobenzyl) glutathione (GS-CNB), (2,4-dinitrobenzyl) glutathione (GS-DNB), imidazole, isopropyl-thio-*β*-D-galactopyranoside (IPTG), lysogeny broth (LB), lysozyme, methanol, methylene chloride, and pepstatin A were the highest grade available from Sigma. Ampicillin, 4-hydroxynonenal (4HNE), KCl, KOH, MgCl_2_, NAD^+^, NADH, and phenylmethylsulfonyl fluoride (PMSF), retinaldehyde, and retinol were purchased from Fisher Scientific. 4-glutathione-nonenal (GS-4HNE) was purchased from Cayman Chemical. Glutathione-, Strep-, and nickel-chelating resins were from GE Life Sciences. Competent *E. coli* (BL21(DE3)) was purchased from Novagen. XL1 Blue competent cells and XL-10 Gold ultracompetent cells were obtained from Agilent. Platinum PCR SuperMix High Fidelity and T4 DNA ligase were obtained from Invitrogen. Gibson Assembly kits and DpnI were purchased from New England Biolabs. The open reading frames of human α- and γ-ADH1, human glutathione S-transferase A1 (GSTA1) and A4 (GSTA4) were obtained from transOMIC Technologies Inc.

### Methods

#### Expression Plasmids

To allow homo- and heterodimers to be separated, Strep- and His-tags (8- and 6-amino acid tags, respectively) were attached to the N-termini of the ADH1 and GST coding regions using Gibson Assembly^57,58^ (Strep-tag, α-ADH1 and GSTA1; His-tag. γ-ADH1 and GSTA4). The tagged coding regions were inserted into a pGEX-6P expression vector containing a PreScission-protease-cleavable N-terminal His/GST/MBP-tag^59,60^. Coding regions were inserted in vectors using Gibson Assembly^57,58^ strategies. Coding region sequences were confirmed *via* DNA sequencing.

#### ADH1 and GST Expression and Purification

*E. coli* (BL21(DE3)) transformed with expression plasmid was grown at 37 °C to an OD_600_ ~ 0.6 in LB/ampicillin (100 mg/L). IPTG (0.30 mM, final) was then added, and the cultures were shifted to 17 °C using an ice/water bath and incubated at 17 °C for 18 hours. Cells were then pelleted, suspended in Lysis Buffer (PMSF (290 µM), pepstain A (1.5 µM), lysozyme (0.10 mg/ml), EDTA (2.0 mM), KCl (400 mM), KPO_4_ (50 mM), pH 7.5) and stirred gently for 1.0 hr at 4 °C. The solution was then sonicated and centrifuged (10,000 g, 1.0 hr) at 4 °C. MgCl_2_ (5.0 mM) was added to the supernatant to sequester EDTA and the solution was passed through a His-column (Chelating Sepharose Fast Flow resin) charged with Ni^2+^ (His-resin). The column was washed with 15 column volumes of Wash Buffer (imidazole (10 mM), KCl (0.40 M), and KPO_4_ (50 mM), pH 7.5, 4 °C). Enzyme was then eluted from the His-column with Elution Buffer (imidazole (250 mM), KCl (0.40 M), KPO_4_ (50 mM), pH 7.5, 4 °C) and loaded directly onto a Glutathione Sepharose column (GST-column) equilibrated with DTT (2.0 mM), KCl (0.40 M), and KPO_4_ (50 mM), pH 7.5, 4 °C. The GST-column was washed with equilibration buffer before eluting the tagged enzyme with reduced glutathione (10 mM), DTT (2.0 mM), KCl (0.40 M), Tris (100 mM), pH 8.0 4 °C). The fusion protein was digested using PreScission Protease during overnight dialysis against: DTT (2.0 mM), KCl (0.10 M), and KPO_4_ (25 mM), pH 7.5, 4°C. To remove tags ADHs were passed through a GST-column; GSTs were passed through a His-column. Cleavage with PreScission protease leaves a GGS-sequence at the N-terminus of the expressed protein. The enzymes are > 95% pure based on SDS-PAGE. Protein concentrations were determined by absorbance using published extinction coefficients^61-63^, aliquoted, flash frozen and stored at −80 °C.

#### ADH1 Heterodimerization

As previously reported^23^, ADH1α and ADH1γ homodimers were monomerized using multiple freeze/thaw cycles and reconstituted as a mixture of homo- and heterodimers after gentle overnight dialysis. Briefly, Strep-tagged ADH1α was mixed with an equimolar concentration (2.0 mg/mL) of His-tagged ADH1γ in buffer (DTT (2.0 mM), glycerol (20%, v/v), NaPO_4_ (500 mM), pH 7.5, 4 °C). The solution was frozen rapidly in dry ice/ethanol, stored overnight at −80 ºC, and thawed at 25 ºC for 2 hr before heterodimerization *via* overnight dialysis against KCl (400 mM), KPO_4_ (50 mM), pH 7.5, 4 °C.

#### GSTA Heterodimerization

GST heterodimers were constructed using the 1,6-hexanediol protocol^64^. Briefly, His-GSTA1 and Strep-GSTA4 homodimers were monomerized separately by incubating the dimers for two hours in 1,6-hexanediol (25% v/v), K_2_PO_4_ (100 mM), EDTA (1.0 mM), pH 7.5, 25 (± 2) °C. Heterodimers were then formed by overnight dialysis of a 1:1 stoichiometric mixture of monomerized GSTA1 and A4 in Tris (10 mM), KCl (200 mM), pH 7.8 at 4°C. The mixture (~ 2ml) was dialyzed three times against 1-liter buffer changes.

#### Separation of Homo- and Heterodimers

Strep-(α-ADH1 and GSTA1) and His-Tags (γ-ADH1 and GSTA4) were used to separate homo-from heterodimers. Only heterodimers harbor both His- and Strep-tags; consequently, only they bind to both His- and Strep-resins. Dual Strep-tagged homodimers are removed from dimer mixtures by loading the dialysate described above onto a His-resin and washing the resin with 15 volumes of Buffer A (imidazole (10 mM), KCl (0.40 M), KPO_4_ (50 mM), pH 7.5, 4 °C). The adsorbed His-tagged dimers (homo- and heterodimers) were then eluted with 8 column volumes of Buffer B (imidazole (250 mM), KCl (0.20 M), K_2_PO_4_ (50 mM), pH 7.5, 4 °C). His-tagged homodimers are removed by loading the eluent directly onto a Strep-Tactin SuperFlow resin and washing the resin with 10 column volumes of Buffer C (Tris (100 mM), KCl (150 mM), pH 8.0, 4 °C). The heterodimer was then eluted with desthiobiotin (2.5 mM in Buffer C), quantitated using the Bradford Assay^65^ and checked for catalytic activity. The heterodimerization and purification protocols yielded heterodimers (ADH1αγ and GSTA1/A4) in were 40-50 % yields.

To ensure heterocomplex did not reformed upon storage at −80 °C, Strep-Tactin resin (sufficient to bind 1.0 mg of protein) was added to 0.10 mg of thawed heterodimer immediately prior to experimentation. The mixture was incubated at 4.0 ºC for one hour, spun gently (1,000 g, 5.0 min, 4 ºC) and the supernatant was assayed. Activity was not detected in either (ADH1 or GSTA) purified heterodimer.

##### Affinity Tags Do Not Effect Activity

The activity of tagged ADH1 (heterodimer and both homodimers) was evaluated using ethanol oxidation. A detailed protocol is presented in “Initial-Rate Studies” and kinetic constants are presented in Table 1. All constants are consistent with literature values for the heterodimer and homodimers^66^. GST-tagged isoforms were evaluated using a common GST assay, CDNB. CDNB assays were performed as DCNB assay described in “Initial-Rate Studies”, with CDNB. Conditions were as follows: GST (5.0 µM, active sites), CDNB (0.20 – 15 mM, 0.20 – 5.0 x K_m_)), GSH (0.20 – 5.0 mM), DTT (2.0 mM), KCl (100 mM), KPO_4_ (50 mM), pH 7.4, 25 ± 2 °C. Activity was detected *via* formation GS-DNB (ε_345_ = 9.6 mM^−1^ cm^−1^). The initial-rate kinetic constants were as follows: GSTA1/A1: K_m CDNB_ = 0.85 ± 0.02 mM, K_m GSH_ = 85 ± 3.3 µM, k_cat_ = 40 ± 3; GSTA4/A4: K_m CDNB_ = 3.6 ± 0.3 mM, K_m GSH_ = 88 ± 2.7 µM, k_cat_ = 33 ± 2.6; GSTA1/A4: K_m CDNB_ = 3.2 ± 0.2 mM, K_m GSH_ = 74 ± 1.7 µM, k_cat_ = 43 ± 2.8. These results agree well with the published literature^17^.

### Coupled-Reaction Equilibrium Studies

#### ADH

The retinol and ethanol reactions were coupled using the α,γ-ADH1 heterodimer. The EOH redox reaction was initiated by addition of NAD^+^ and reaction progress was monitored continuously *via* NADH absorbance at 340 nM. The EOH reaction was allowed to reach equilibrium before adding ROH, at 60 min, to 4.0 µM. The ROH reaction was monitored continuously *via* RA fluorescence (λ_ex_ = 420 nm, λ_em_ = 530 nm). AA was calculated as: [NADH] – [RA]. Reaction conditions at t_o_: α,γ-ADH1 (0.10 nM, heterodimer), NAD^+^ (10 mM, 25 x K_m_), ethanol (4.0 µM), KPO_4_ (50 mM), KCl (100 mM) pH 7.4, 25 ± 2ºC.

#### GST

The DCNB and 4HNE reactions were coupled using the GSTA1/A4 heterodimer. Conditions were as follows: GSTA1/A4 heterodimer or a 1:1 mixture of GSTA1 and GSTA4 homodimers (100 nM, dimers), DCNB (100 µM, 59 x K_m_), 4HNE (200 µM, 4.4 x K_m_), GSH (300 µM, 4.0 x K_m_), DTT (2.0 mM), and KPO_4_ (50 mM, pH 7.4). DCNB and GS-CNB were monitored *via* absorbance (ε_345_ = 8500 M^−1^ cm^−1^) and fluorescence (λ_ex_ = 325 nm and λ_em_ = 375 nm), respectively. 4HNE and GS-HNE (black) were monitored *via* absorbance (ε_225_ = 19200 M^−1^ cm^−1^ and ε_285_ = 19500 M^−1^ cm^−1^, respectively). Each progress curve is the average of three reactions and the line through the points is predicted by the model (see, *Supplement 1, Modelling*) using the kinetic constants in Table 4.

#### GS-4HNE ^1^H-NMR

Conditions: GS-4HNE (2.0 mM), K_2_PO_4_ (50 mM), pH 7.4 25 ± 2 °C. D_2_O (> 99%). TSP (0.20 mM) was present in a coaxial NMR sample-tube insert. Spectra were acquired with water suppression using a Bruker 600 MHz spectrometer equipped with a TCI H/F-cryogenic probe ^67^. Each spectrum was the average of 1024 scans acquired using a 1D-pulse sequence (20.0 s relaxation delay, s acquisition time). The relaxation delay used in 1D-NMR studies was ≥ 5.0 times T1 of the cyclic proton (i.e., the proton with the longer T1, 3.2 s). T1 values were determined using a standard T1 *inversion*-recovery protocol ^68^. Peaks were fit to a Lorentzian shape using TopSpin3.5 ^42^ and concentrations were determined by normalizing peak areas to that of the TSP signal. Assignment the cyclic H1 resonance was based on published spectra^41^. The linear H1 resonance was assigned based on its predicted chemical shift and saturation transfer between the linear and cyclic resonances.

## Abbreviations

ADH: (alcohol dehydrogenase)
DCNB: (1,2-dichloro-4-nitrobenzene)
GS-CNB: 1-(glutathion-S-yl)-2-chloro-4-nitrobenzene)
GS-HNE: (glutathionyl-4-hydroxynonanal)
GSH: (glutathione)
GST: (glutathione S-transferases)
4-HNE: (4-hydroxy-2-nonenal)
RA: (retinaldehyde)
ROH: (retinol)

## Acknowledgements

This work was supported by National Institutes of Health Grants GM121849 and GM127144 to TSL.

We thank Dr. Sean Cahill for his excellent guidance regarding the NMR experimentation described herein. The Bruker 600 MHz NMR instrument in the Einstein Structural NMR Resource was purchased using funds from NIH award 1S10OD016305 and is supported by the Albert Einstein College of Medicine.

## Author Contributions

I.C. and T.W performed all experiments, I.C. assisted T.S.L in experimental design and interpretation, and in writing and assembling the manuscript. T.S.L. conceived the project, guided experimentation and interpretation, and wrote the majority of the manuscript.

