## Supplementary Materials for "On the Distribution of Free-Energy in Metabolism"

#### *On the Energetics of Small-Molecule Metabolism*

**Supplemental materials include:**

Figs. S1 to S13

***Initial-Rate Studies.***

***Overview.*** Experimental conditions associated with the ADH1 and GSTA initial-rate studies are detailed in the corresponding figure legends. Except for the two cases discussed below (see, *Substrate-Competition Studies*) all studies were performed at a fixed-saturating concentration (12 - 30 $K_m$) of one or more substrates and the variable-substrate concentration ranged from ~ 0.20 $K_m$ to 5.0 $K_m$ in equal increments in double-reciprocal space. In all cases, consumption of the concentration-limiting substrate was < 5% of product formed at the reaction endpoint. All rate measurements were performed in triplicate. Unless otherwise stated, initial-rate parameters were estimated using *sequeno*, a $1/v^4$-weighted least-squares fitting algorithm[4-6].

ADH1 reaction progress was monitored by detecting formation of one of four reactants: **1**, NADH, monitored continuously *via* NADH absorbance ($\varepsilon_{340}$ = 6.22 $mM^{-1}$ $cm^{-1}$); **2**, $NAD^+$, detected by optically quantitating (260 nm) dinucleotides separated using HPLC (see, *HPLC-Based Detection*); **3**, ROH, monitored *via* a gain in ROH fluorescence ($\lambda_{ex}$ = 370 nm, $\lambda_{em}$ = 450 nm); **4**, RA, detected by a gain in RA fluorescence ($\lambda_{ex}$ = 420 nm, $\lambda_{em}$ = 550 nm).

GST reaction progress was monitored by detecting formation of one of four reactants: **1**, GS-CNB, fluorescence gain ($\lambda_{ex}$ = 345 nm and $\lambda_{em}$ = 375 nm); **2**, DCNB, absorbance gain ($\varepsilon_{315}$ = 8,500 $M^{-1}$ $cm^{-1}$); **3**, GS-4HNE, absorbance loss ($\varepsilon_{225}$ = 19,500 $M^{-1}$ $cm^{-1}$); and **4**, GSH, gain in absorbance of 2-nitro-5-mercapto-benzoic acid TNB ($\lambda_{ex}$ = 412 nm, $\lambda_{em}$ = 550 nm) produced by non-enzymatic conversion of GSH to TNB by reaction with DTNB (5.0 mM) [8]. All reactions were monitored continuously.

*DCNB Studies*. The literature reports that DCNB is not a substrate for the GSTA4 homodimer[9]. To confirm this assertion, the activity of the GSTA4 homodimer toward DCNB was assessed. Reaction progress was monitored *via* GS-CNB fluorescence ($\lambda_{ex}$ = 345 nm and $\lambda_{em}$ = 375 nm) under the following conditions: GSTA4 (25 μM), DCNB (1.0 mM), GSH (5.0 mM), $KPO_4$ (50 mM), KCl (100 mM), pH 7.4, 25 ± 2 °C. The studies (repeated in triplicate) showed no detectible GS-CNB formation over 1 hr. Given a lower limit of GS-CNB detection of ~100 nM, GSTA4 turnover of DCNB is calculated at ≤ 0.004 $h^{-1}$, which is too slow to meaningfully contribute to our GST initial-rate and equilibrium studies.

To assess whether DCNB binds the GSTA4 active site of the heterodimer and thus competes with 4HNE in the coupled reaction studies. DCNB was tested as an inhibitor of the 4HNE reaction under the following conditions: DCNB (1.0 mM), 4HNE (1.0 mM, 3,600 x $K_m$), GSH (5.0 mM, 45 x $K_m$), GSTA4 (25 μM), $KPO_4$ (50 mM), KCl (100 mM), pH 7.4, 25 ± 2 °C. DCNB did not detectibly inhibit the 4HNE reaction at 1.0 mM, which is 6.6-fold higher than the highest DCNB concentration used in either the hetero- or homodimer studies.

*Substrate-Competition Studies*. In the majority of heterodimer coupled-reaction initial-rate studies, it was possible, due to orders-of-magnitude differences between coupled and uncoupled reactant $K_m$ and $k_{cat}$ values, to choose conditions such that turnover of the monitored reaction occurred nearly exclusively (> 97%) through the heterocomplex reaction pathway. This was not the case for three studies (*1/v-vs-1/AA,* Fig. S3G; *1/v-vs-1/ROH*, Fig. S4E; and *1/v-vs-1/DCNB,* Fig. S7B) either because substrate $K_m$ values are comparable for both subunits (S3G and S4E), or the homocomplex $k_{cat}$ is greater than that of the heterocomplex (S7B, Table 4). Given $K_{m\,S1}$, obtained from heterodimer single-reaction studies, both $K_{m\,S2}$ and $k_{cat}$ for the heterocomplex can be obtained in a substrate competition study in which the S2 reaction rate is monitored as a function of both S1 and S2 concentrations. Competition studies between AA and RA, ROH and EOH, and DCNB and 4HNE were performed using 4 x 4 substrate-concentration matrices. Reaction conditions are described in the Fig S3, S4, and S7 figure legends. Initial-rate parameters were obtained by global least-squares fitting using the following competition model**:** $\nu_A = (A) \bullet V_{max\,A} / [K_{m\,A} (1 + (B) / K_{m\,B}) + A]$, where $\nu_A$ is the rate at which A is consumed, (A) and (B) are substrate concentrations. $V_{max\,A}$ and $K_{m\,A}$ were obtained by fitting the 4 x 4 at a fixed value of $K_{m\,B}$, which was obtained from single-reaction studies.

***ADH1-Specific Initial-Rate Protocols.***

*HPLC Based Detection***.** The $NAD^+$ produced in the acetaldehyde reduction studies was too small relative to NADH levels for continuous UV monitoring; consequently, $NAD^+$ and NADH were separated using HPLC and quantitated. Reactions (25 µl) were stopped after 1.0 – 10 min by addition of NaOH (0.10 M, final), neutralized 1 min later by adding HCl and then diluted by addition of 25 µl of $NaPO_4$ (50 mM), pH 7.0. The quenched solutions were filtered (0.22 µm) and the filtrate was loaded onto a VYDAC™ C18-column. NADH and $NAD^+$ were quantitated (260 nm) by indexing HPLC peak areas to standard curves constructed using commercial compounds. Duplicate four-point progress curves were constructed under each set of conditions. The data were averaged and velocities were obtained by linear least-squares fitting. Chromatographic separation of $NAD^+$ and NADH was achieved as follows: *Buffer A*, 0.10 M sodium dihydrogen phosphate, 8.0 mM tetrabutylammonium hydrogen sulfate, pH 6.0; *Buffer B*, 0.10 M sodium dihydrogen phosphate, 8.0 mM tetrabutylammonium hydrogen sulfate, 30% methanol, pH 6.0; flow rate, 1.0 ml/min; gradient: 0', 100% A; 4', 100% A; 8', 80% A; 16', 60% A; 20', 100% A; 28', 0% A; 30', 100% A.

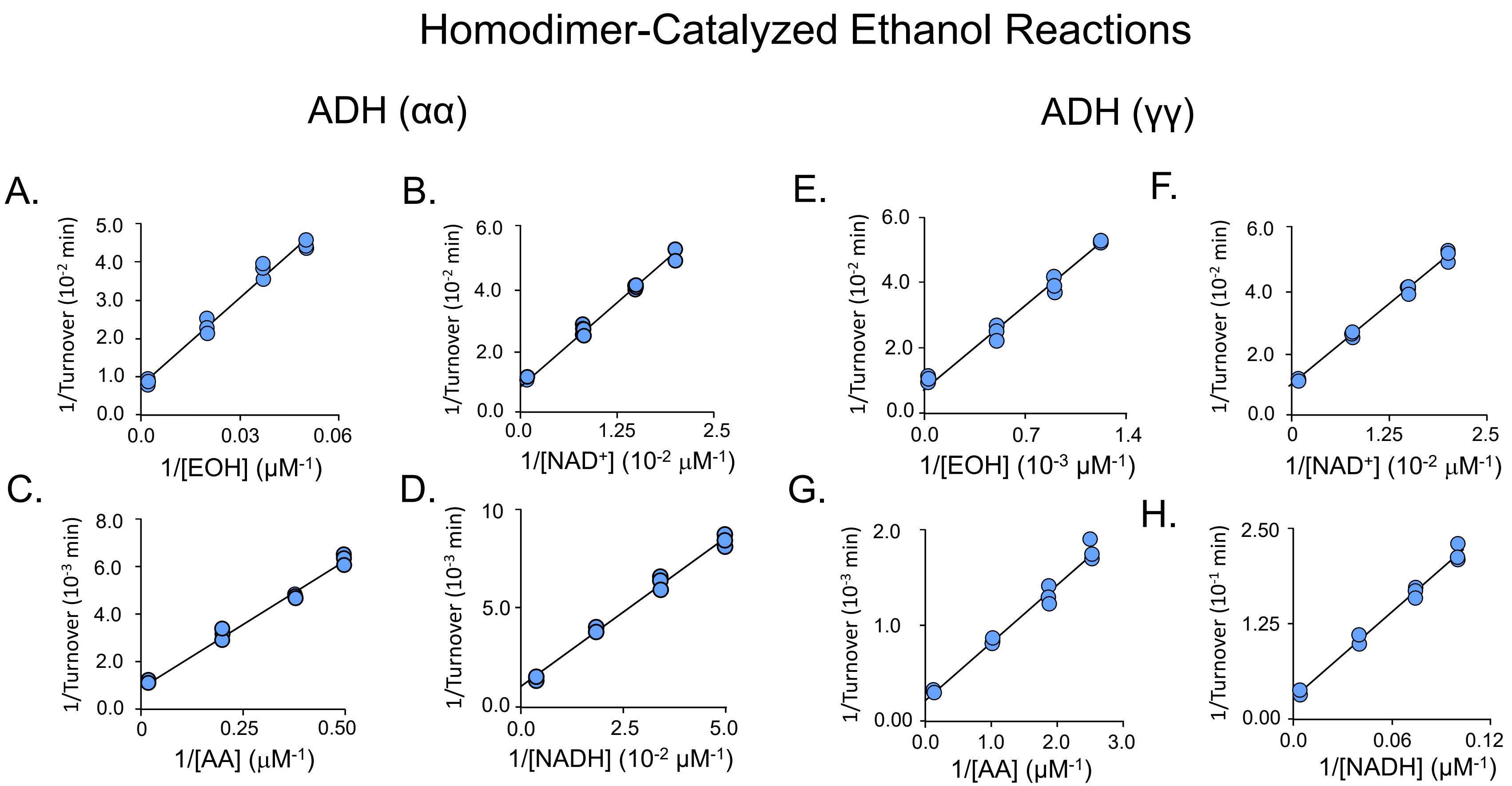


**Fig. S1. Initial-rate studies of the ethanol redox reactions catalyzed by αα-ADH1 and γγ-ADH1.** *αα-ADH1(A-D) and γγ-ADH1(E-H) studies.* The x-axis variable-substrate concentrations in each study ranged from ~ 0.20 – 5.0 x $K_m$ in equal increments in double-reciprocal space. The following conditions were used in all studies: $KPO_4$ (50 mM), KCl (100 mM), pH 7.4, 25 ± 2 °C. **(A)** and **(E)** *1/Turnover vs 1/[EOH]*. Conditions: ADH1 (50 nM active sites), [$NAD^+$] (6.0 mM, 19 and 29 x $K_m$ for αα- and γγ-ADH1, respectively). **(B)** and **(F)** *1/Turnover vs 1/[$NAD^+$]*. Conditions: ADH1 (50 nM active sites), [EOH] (100 mM, 27 and 830 x $K_m$ for αα- and γγ-ADH1, respectively). **(C)** and **(G)** *1/Turnover vs 1/[AA]*. Conditions: ADH1 (10 nM active sites), NADH (4.0 mM, 59 and 310 x $K_m$ for αα- and γγ-ADH1, respectively). (**D**) and **(H)** *1/Turnover vs 1/[NADH]*. Conditions: ADH1 (10 nM active sites), [AA] (5.0 mM, 19 and 420 x $K_m$ for αα- and γγ-ADH1, respectively). Forward and reverse reactions were monitored *via* NADH and $NAD^+$ absorbance, respectively (see, *Supplementary Materials*, *ADH Initial-Rate Studies*). Data were fit using $1/v^4$-weighted, linear least-squares analysis and the resulting best-fit initial-rate constants are compiled in Table 1

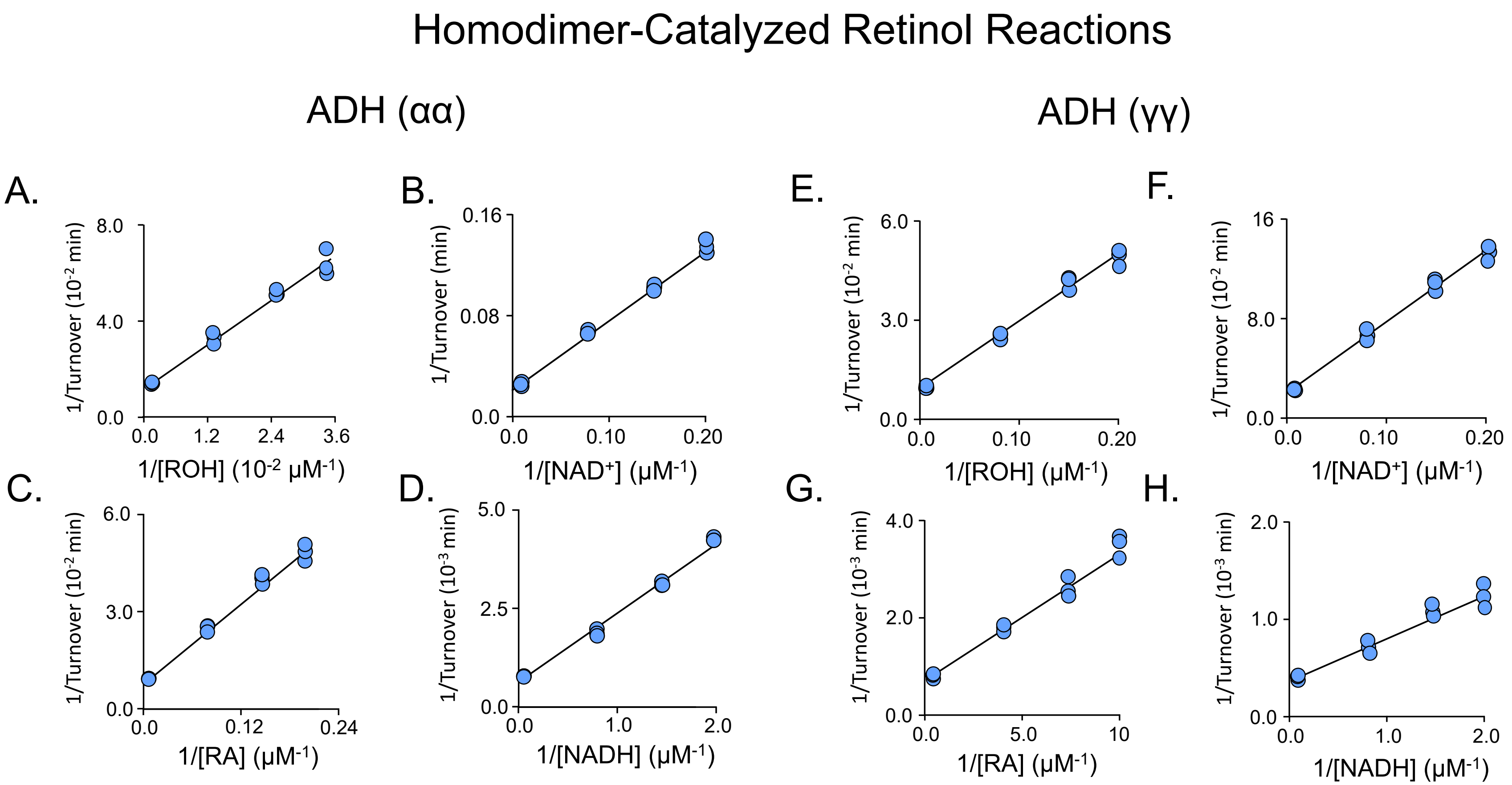


**Fig. S2. Initial-rate studies of the retinol redox reactions catalyzed by αα-ADH1 and γγ-ADH1.** *αα-ADH1 (A-D) and γγ-ADH1 (E-H) studies.* The x-axis variable-substrate concentrations in each study ranged from ~0.20 – 5.0 x $K_m$ in equal increments in double-reciprocal space. The following conditions were used in all studies: $KPO_4$ (50 mM), KCl (100 mM), pH 7.4, 25 ± 2 °C. **(A)** and **(E)** *1/Turnover vs 1/[ROH].* Conditions: ADH1 (100 nM active sites), [$NAD^+$] (6.0 mM, 19 and 26 x $K_m$ for αα- and γγ-ADH1, respectively). **(B)** and **(F)** *1/Turnover vs 1/[$NAD^+$].* Conditions: ADH1 (100 nM active sites), [ROH] (5.0 mM, 111 and 26 x $K_m$ for αα- and γγ-ADH1, respectively). **(C)** and **(G)** *1/Turnover vs 1/[RA].* Conditions: ADH1 (20 nM active sites), NADH (50 μM, 46 and 20 x $K_m$ for αα- and γγ-ADH1, respectively). (**D**) and **(H)** *1/Turnover vs 1/[NADH].* Conditions: ADH1 (20 nM active sites), [RA] (15 μM, 33 and 32 x $K_m$ for αα- and γγ-ADH1, respectively). Forward and reverse reactions were monitored *via* RA or ROH fluorescence, respectively (see, *Supplementary Materials*, *ADH Initial-Rate Studies*). Data were fit using $1/v^4$-weighted, linear least-squares analysis and the resulting best-fit initial-rate constants are compiled in Table 1

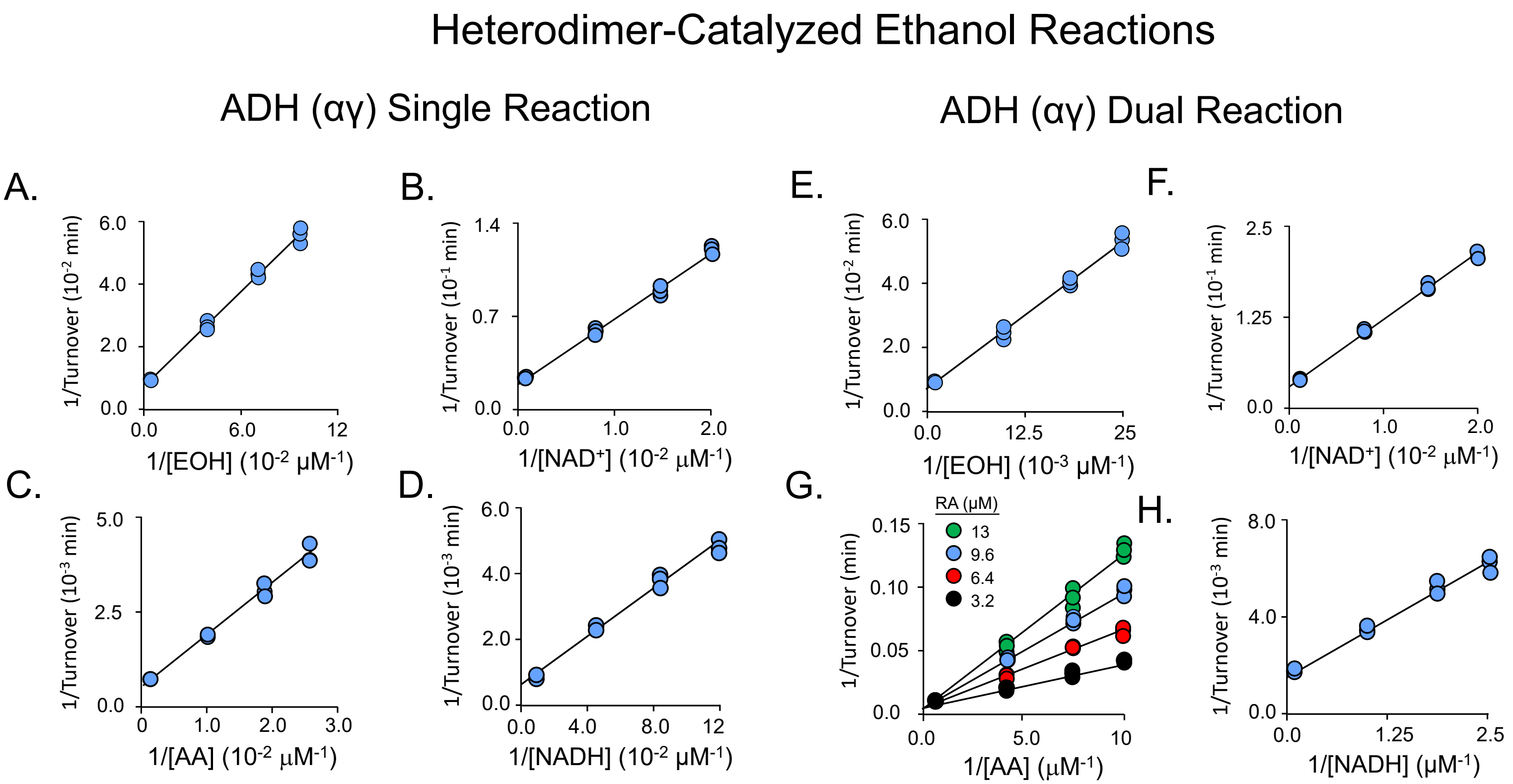


**Fig. S3. Initial-rate studies of the uncoupled and retinol-coupled ethanol redox reactions catalyzed by the αγ-ADH1 heterodimer.** *Uncoupled (A-D) and coupled (E-H) studies.* The variable-substrate concentrations in each study ranged from ~0.20 – 5.0 x $K_m$ in equal increments in double-reciprocal space. The following conditions were used in all studies: $KPO_4$ (50 mM), KCl (100 mM), pH 7.4, 25 ± 2 °C. **(A)** *1/Turnover vs 1/[EOH].* Conditions: *αγ*-ADH1 (50 nM active sites), [$NAD^+$] (6.0 mM, 27 x $K_m$). **(B)** *1/Turnover vs 1/[$NAD^+$].* Conditions: *αγ*-ADH1 (50 nM active sites), [EOH] (100 mM, 30 x $K_m$). **(C)** *1/Turnover vs 1/[AA].* Conditions: *αγ*-ADH1 (10 nM active site), NADH (1.5 mM, 23 x $K_m$). (**D**) *1/Turnover vs 1/[NADH].* Conditions: *αγ*-ADH1 (10 nM active sites), [AA] (5.0 mM, 24 x $K_m$). **(E)** *1/Turnover vs 1/[EOH] at saturating [ROH] and [$NAD^+$].* Conditions: *αγ*-ADH1 (10 nM active site), ROH (5.0 μM, 26 x $K_m$), $NAD^+$ (6.0 mM, 16 x $K_m$). **(F)** *1/Turnover vs 1/[$NAD^+$] at saturating [ROH] and [EOH].* Conditions: *αγ*-ADH1 (10 nM active site), ROH (5.0 μM, 26 x $K_m$), [EOH] (5.0 mM, 23 x $K_m$). **(G)** *1/Turnover vs 1/[AA] at fixed-variable [RA].* Conditions: *αγ*-ADH1 (10 nM active site), NADH (30 μM, 30 x $K_m$). Due to the inability to achieve the desired subunit ligand distributions at a single [RA], a substrate competition experiment was performed (see, *Supplemental*, *Initial-Rate Studies*). Rates were determined at each of the 16 conditions defined by a 4 x 4 concentration matrix in which AA varied from 0.25 – 10 x $K_m$ (0.10 – 10 μM) and RA varied from 2.5 – 20 x $K_m$ (3.2 – 13 μM) — [S]/$K_m$ values are calculated based on $K_m$ for the lower-affinity subunit. The high-affinity subunit was near saturation (7.1 – 37 x $K_m$) at all [RA]. **(H)** *1/Turnover vs 1/[NADH] at saturating [RA] and [AA].* Conditions: ADH (5.0 nM active site), RA (10 μM, 29 x $K_m$), AA (5.0 μM, 12 x $K_m$). Progress of the forward and reverse reactions was monitored *via* NADH and $NAD^+$ absorbance, respectively (see, *Supplementary Materials*, *ADH Initial-Rate Studies*). Data were fit using $1/v^4$-weighted, linear least-squares analysis and the resulting best-fit initial-rate constants are compiled in Table 2.

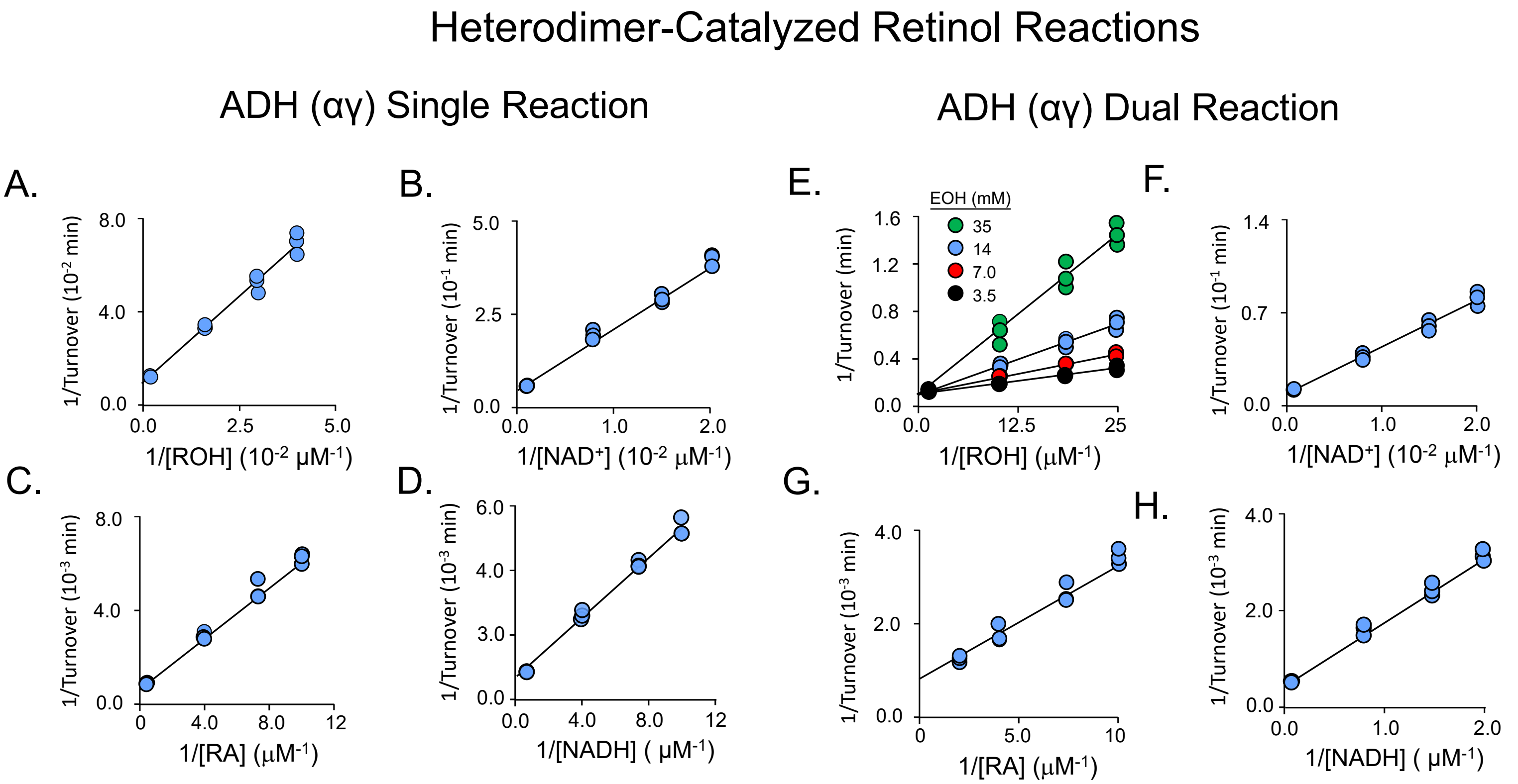


**Fig. S4. Initial-rate studies of the uncoupled and ethanol-coupled retinol redox reactions catalyzed by the αγ-ADH1 heterodimer.** *Uncoupled (A-D) and coupled (E-H) studies.* The variable-substrate concentrations in each study ranged from ~0.20 – 5.0 x $K_m$ in equal increments in double-reciprocal space. The following conditions were used in all studies: $KPO_4$ (50 mM), KCl (100 mM), pH 7.4, 25 ± 2 °C. **(A)** *1/Turnover vs 1/[ROH].* Conditions: *αγ*-ADH1 (50 nM active sites), [$NAD^+$] (6.0 mM, 15 x $K_m$). **(B)** *1/Turnover vs 1/[NAD$^+$].* Conditions: *αγ*-ADH1 (50 nM active sites), [ROH] (3.5 μM, 18 x $K_m$). **(C)** *1/Turnover vs 1/[RA].* Reaction conditions: *αγ*-ADH1 (10 nM active site), NADH (30 μM, 25 x $K_m$). (**D**) *1/Turnover vs 1/[NADH].* Conditions: *αγ*-ADH1 (10 nM active site), [RA] (15 μM, 43 x $K_m$). **(E)** *1/Turnover vs 1/[ROH] at fixed-variable [EOH].* Conditions: ADH (50 nM active site), [$NAD^+$] (6.0 mM, 21 x $K_m$). Due to the inability to achieve the desired subunit ligand distributions at a single [EOH], a substrate competition experiment was performed (see, *Supplemental*, *Initial-Rate Studies*). Rates were determined at each of the 16 conditions defined by a 4 x 4 concentration matrix in which ROH varied from 0.20 – 5.0 x $K_m$ (0.040 – 1.0 μM) and EOH varied from 1.1 – 11 x $K_m$ (3.5 – 35 mM) of the low-affinity subunit. The heterodimer subunit with high affinity for ETOH subunit was saturated (15 - 160 x $K_m$) at all [EOH]. **(F)** *1/Turnover vs 1/[NAD$^+$] at saturating [EOH] and [ROH].* Conditions: *αγ*-ADH1 (2.0 nM active site), EOH (5.0 mM, 23 x $K_m$), ROH (6.0 μM, 31 x $K_m$). **(G)** *1/Turnover vs 1/ [RA] at saturating [AA] and [NADH].* Conditions: *αγ*-ADH1 (10 nM active site), AA (5.0 μM, 12 x $K_m$), NADH (30 μM, 23 x $K_m$). (**H**) *1/Turnover vs 1/[NADH] at saturating [RA] and [AA].* Conditions: *αγ*-ADH1 (5.0 nM active site), RA (10 μM, 29 x $K_m$), AA (5.0 μM, 12 x $K_m$). Forward and reverse reactions were monitored *via* an increase in RA or ROH fluorescence ($\lambda_{ex\ ROH}$ = 370 nm, $\lambda_{em\ ROH}$ = 450 nm; $\lambda_{ex\ RA}$ = 420 nm, $\lambda_{em\ RA}$ = 550 nm). Data were fit using $1/v^4$-weighted, linear least-squares analysis and the resulting best-fit initial-rate constants are compiled in Table 2.

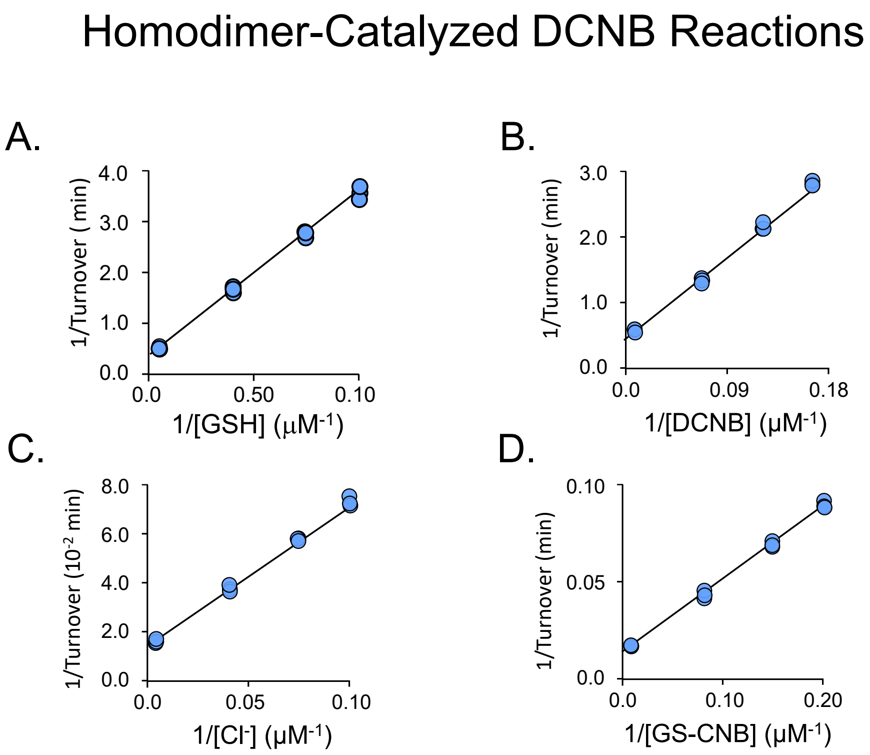


**Fig. S5. Initial-rate studies of DCNB/GSH conjugation reaction catalyzed by GSTA1.** The variable-substrate concentrations in each study ranged from ~0.20 – 5.0 x $K_m$ in equal increments in double-reciprocal space. The following conditions apply to all studies: $KPO_4$ (50 mM), pH 7.4, 25 ± 2 °C. The fixed substrate concentration was saturating in all cases (> 15 x $K_m$). **(A)** *1/Turnover vs 1/[GSH].* Conditions: GSTA1 (10 nM, dimer), [DCNB] (600 μM 17 x $K_m$). **(B)** *1/Turnover vs 1/[DCNB].* Conditions: GSTA1 (10 nM, dimer), GSH (3.5 mM, 45 x $K_m$). Forward reactions were monitored continuously *via* GS-CNB fluorescence ($\lambda_{ex}$ = 345 nm and $\lambda_{em}$ = 375 nm). **(C)** *1/Turnover vs 1/[$Cl^-$].* Conditions: GSTA1 (25 nM, dimer), GS-CNB (1.5 mM, 60 x $K_m$). The $Cl^-$ concentration includes buffer-background [$Cl^-$] (6.1 μM, see *DNCB Conjugation* in *Methods*). (**D**) *1/Turnover vs 1/[GS-CNB].* Conditions: GSTA1 (25 nM, dimer), $Cl^-$ (2.0 mM, 50 x $K_m$). Reverse reactions were monitored continuously *via* DCNB absorbance ($\varepsilon_{315}$ = 8,500mM$^{-1}$ cm$^{-1}$). In all studies, consumption of the concentration-limiting substrate was ≤ 5% of product formed at the reaction endpoint. Each measurement was performed in triplicate. Data were fit using 1/v$^4$-weighted, linear least-squares analysis and the resulting best-fit initial-rate constants are compiled in Table 3. GSTA4 activity was not detected (≤ 0.004 turnover) over 1.0 h at: GSTA4 (25 μM), DCNB (1.0 mM), GSH (5.0 mM).

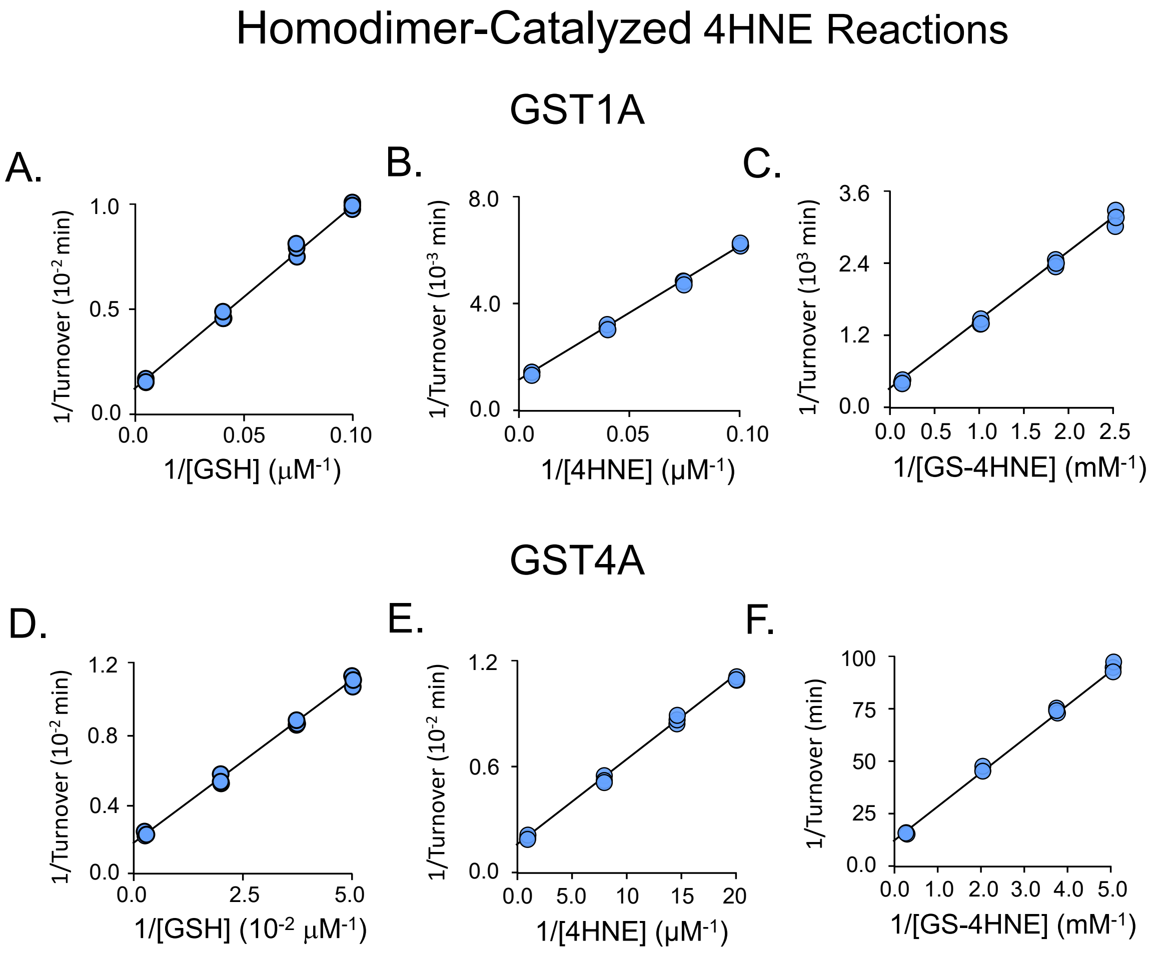


**Fig. S6. Initial-rate studies of the uncoupled 4HNE GSH-conjugation reactions catalyzed by GST homodimers.** *GSTA1 (A-C) and GSTA4 (D-F) studies***.** The variable-substrate concentrations in each study ranged from ~0.20 – 5.0 x $K_m$ in equal increments in double-reciprocal space. The following condition applies to all studies: $KPO_4$ (50 mM), pH 7.4, 25 ± 2 °C. **(A)** and **(D),** *1/Turnover vs 1/[GSH].* Conditions: GST (5.0 nM for GSTA1, 1.0 nM for GSTA4), 4HNE (800 μM, 20 and 2900 x $K_m$ for GSTA1 and GSTA4, respectively). **(B)** and **(E)** *1/Turnover vs 1/[4HNE].* Conditions: GST (50 nM for GSTA1, 5.0 nM for GSTA4), GSH (3.5 mM, 48, 32 x $K_m$ for GSTA1 and GSTA4, respectively). Forward reactions were monitored continuously *via* GS-4HNE absorbance ($\varepsilon_{225}$ = 19,500 $M^{-1}$ $cm^{-1}$). **(C)** and **(F)** *1/Turnover vs 1/[GS-4HNE].* Conditions: GST (10 nM active site) and DTNB (5.0 mM). Formation of GSH was monitored *via* non-enzymatic reaction of GSH with DTNB to form TNB, which is fluorescence ($\lambda_{ex}$ = 412 nm, $\lambda_{em}$ = 540 nm). In all studies, consumption of the concentration-limiting substrate was ≤ 5% of product formed at the reaction endpoint. Each measurement was performed in triplicate. Data were fit using $1/v^4$-weighted, linear least-squares analysis and the resulting best-fit initial-rate constants are compiled in Table 3.

Heterodimer-Catalyzed DCNB Reactions

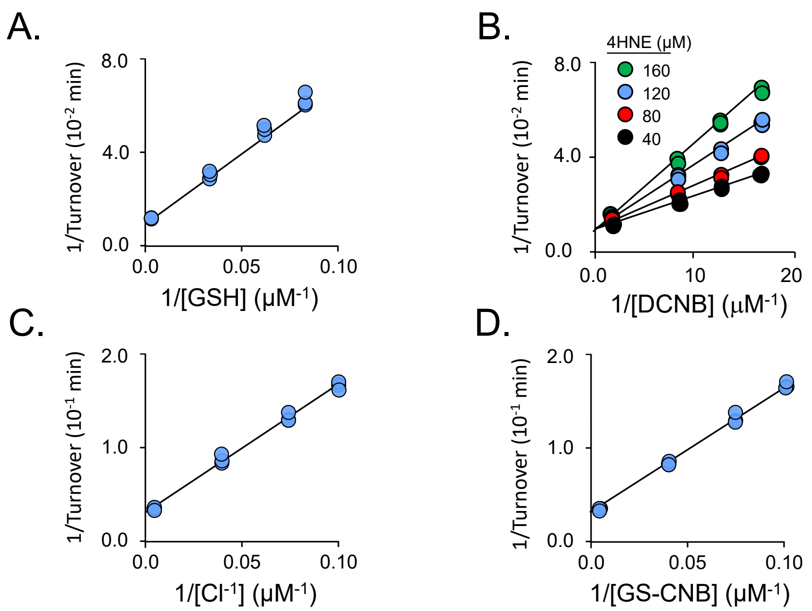


**Fig. S7. Initial-rate studies of the 4HNE-coupled DCNB reactions catalyzed by the GSTA1/A4 heterodimer.** The variable-substrate concentrations in each study ranged from ~0.20 – 5.0 x $K_m$ in equal increments in double-reciprocal space. The following condition applies to all studies: $KPO_4$ (50 mM), pH 7.4, 25 ± 2 °C. **(A)** *1/Turnover vs 1/[GSH] at saturating [DCNB] and [4HNE].* Conditions: GSTA4/A1 (25 nM, dimer), DCNB (50 µM, 500 x $K_m$), 4HNE (200 µM, 17 and 4.5 x $K_m$ for the A4 and A1 subunit, respectively). The [GSH] varied from 10 - 250 µM (0.1 – 3.5 x $K_m$) and reactions were monitored *via* GS-CNB fluorescence ($\lambda_{ex}$ = 345 nm, $\lambda_{em}$ = 375 nm). **(B)** *1/Turnover vs 1/[DCNB] at fixed-variable [4HNE].* Conditions: GSTA4/A1 (5.0 nM, dimers) and GSH (1.0 mM, 17 x $K_m$). Due to the inability to achieve the desired subunit ligand distributions at a single [4HNE], a substrate competition experiment was performed (see, *Supplemental*, *Initial-Rate Studies*). Rates were determined at each of the 16 conditions defined by a 4 x 4 concentration matrix in which DCNB varied from 0.25 – 10 x $K_m$ (20 – 500 nM) and 4HNE varied from 0.91 – 3.6 x $K_m$ (40 – 160 µM) — [S]/$K_m$ values are calculated based on $K_m$ for the lower-affinity subunit. Notably, the heterodimer subunit with higher HNE affinity was at or near saturation (3.8 – 13 x $K_m$) at all 4HNE concentrations, and DCNB conjugation was not detected in the absence of 4HNE (i.e., ≤ 0.004 turnover $h^{-1}$ at GSTA1/A4 (25 µM), DCNB (1.0 mM), GSH (5.0 mM)). **(C)** *1/Turnover vs 1/[$Cl^-$] at saturating [GS-4HNE] and [GS-CNB].* Conditions: GSTA4/A1 (50 nM, dimer), GS-CNB (1.0 mM, 16 x $K_m$), GS-4HNE (4.5 mM, 5.0 and 1.3 x $K_m$ for the A4 and A1 subunit, respectively). $Cl^-$ was varied from 10 – 250 µM (0.22 – 6.1 x $K_m$). The reverse reaction was monitored by DCNB absorbance ($\varepsilon_{315}$ = 8,500 $M^{-1}$ $cm^{-1}$). The $Cl^-$ concentration includes buffer background $Cl^-$ (6.1 µM, see *DCNB Conjugation* in *Methods*). **(D)** *1/Turnover vs 1/[GS-CNB] at saturating [GS-4HNE] and [$Cl^-$].* Conditions: GSTA4/A1 (50 nM, dimer), GS-4HNE (4.5 mM, 5.0 and 1.3 x $K_m$ for the A4 and A1 subunit, respectively), $Cl^-$ (1.0 mM, 22 x $K_m$), GS-CNB was varied from 10 – 250 µM (0.16 – 4.1 x $K_m$)**.** In all studies, consumption of the concentration-limiting substrate was ≤ 5% of product formed at the reaction endpoint. Each measurement was performed in triplicate. Data were fit using $1/v^4$-weighted, linear least-squares analysis and the resulting best-fit initial-rate constants are compiled in Table 4.

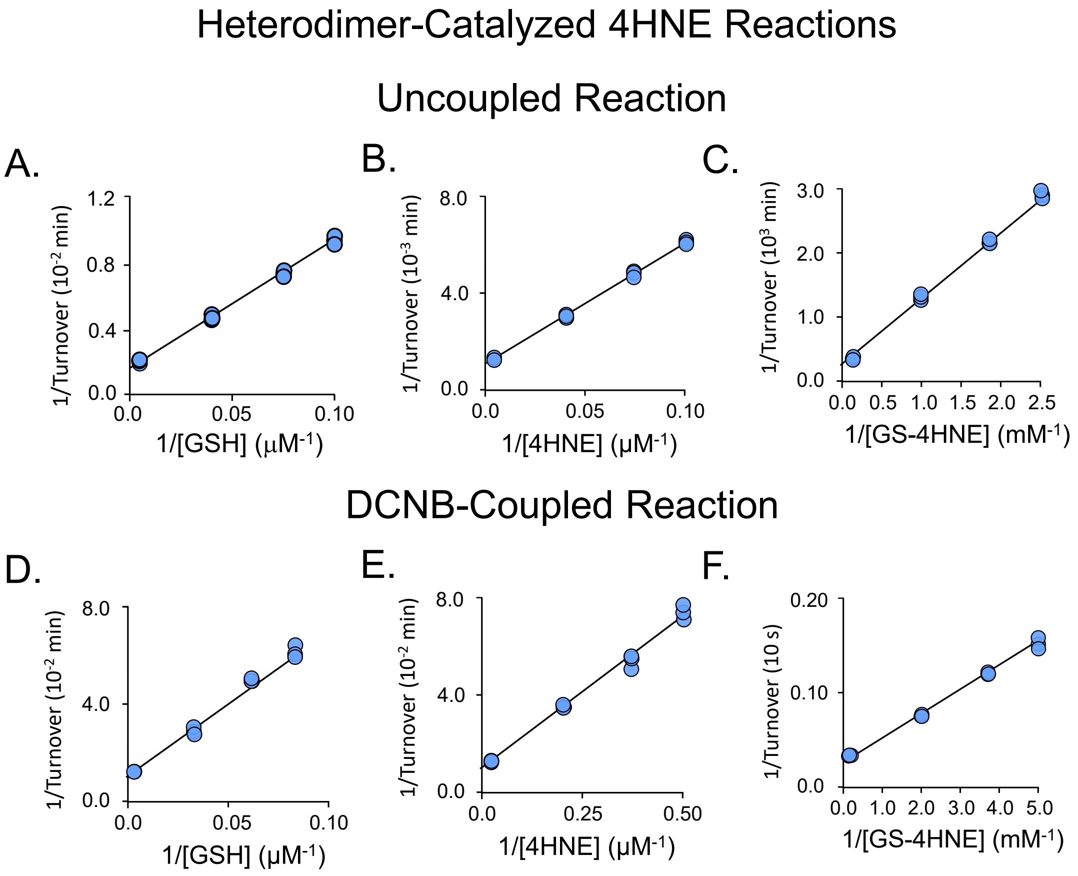


**Fig. S8. Initial-rate studies of the uncoupled and DCNB-coupled 4HNE reactions catalyzed by the GSTA1/A4 heterodimer.** *Uncoupled (A-C) and DCNB-coupled (D-F) GSTA1/A4 studies.* The variable-substrate concentrations in each study ranged from ~0.20 – 5.0 x $K_m$ in equal increments in double-reciprocal space. The following condition applies to all studies: $KPO_4$ (50 mM), pH 7.4, 25 ± 2 °C. **(A)** *1/Turnover vs 1/[GSH].* Conditions: GSTA1/A4 (5.0 nM, dimer), 4HNE (800 μM, 18 x $K_m$). **(B)** *1/Turnover vs 1/[4HNE].* Conditions: GSTA1/A4 (5.0 nM, dimer), GSH (3.5, 49 x $K_m$). Forward reactions were monitored continuously *via* GS-4HNE absorbance ($\varepsilon_{225}$ = 19,500 $M^{-1}$ $cm^{-1}$). **(C)** *1/Turnover vs 1/[GS-4HNE].* Condition: GSTA1/A4 (5.0 nM, dimer), DTNB (5.0 mM). Formation of GSH was monitored *via* the non-enzymatic reaction of GSH with DTNB to form TNB, which is fluorescent ($\lambda_{ex}$ = 412 nm, $\lambda_{em}$ = 540 nm). **(D)** *1/Turnover vs 1/[GSH] at saturating [DCNB] and [4HNE].* Conditions: GSTA1/A4 (25 nM, dimer), DCNB (10 μM, 100 x $K_m$), 4HNE (150 μM, 1.5 and 3.5 x $K_m$ for the A4 and A1 subunit, respectively). GSH was varied from 10 - 250 μM (0.1 – 3.5 x $K_m$). The forward reactions were monitored continuously *via* GS-CNB fluorescence ($\lambda_{ex}$ = 345 nm, $\lambda_{em}$ = 375 nm). **(E)** *1/Turnover vs 1/[4HNE] at saturating [DCNB] and [GSH].* Conditions: GSTA4/A1 (2.5 nM, dimer), DCNB (10 μM, 100 x $K_m$), GSH (1.0 mM, 17 x $K_m$), 4HNE was varied from 2.0 to 50 μM (0.17 – 4.2 x $K_m$). **(F)** *1/Turnover vs 1/[GS-4HNE] at saturating [GS-CNB] and [$Cl^-$].* Conditions: GSTA1/A4 (50 nM, dimer), $Cl^-$ (1.0 mM, 22 x $K_m$), GS-CNB (1.2 mM, 20 x $K_m$), DTNB (5.0 mM), GS-4HNE was varied from 10 – 250 μM (0.16 – 4.1 x $K_m$). Formation of GSH was monitored *via* the non-enzymatic reaction of GSH with DTNB to form TNB, which is fluorescent ($\lambda_{ex}$ = 412 nm, $\lambda_{em}$ = 540 nm). In all studies, consumption of the concentration-limiting substrate was ≤ 5% of product formed at the reaction endpoint. Each measurement was performed in triplicate. Data were fit using $1/v^4$-weighted, linear least-squares analysis and the resulting best-fit initial-rate constants are compiled in Table 4.

##### *Equilibrium Constant Determinations (Single Reaction Studies)*

The following protocols are associated with Fig S9A-D.

##### *ADH*

***EOH Redox (Fig S9A).*** The reaction was monitored *via* NADH absorbance ($\varepsilon_{340}$ = 6.22 $mM^{-1}$ $cm^{-1}$). The $t_0$ condition: γγ-ADH1 (10 μM, active sites), EOH (50 mM), $NAD^+$ (1.0 mM), $KPO_4$ (50 mM), KCl (100 mM), pH 7.4, 25 ± 2 °C. Once the reaction achieved equilibrium, the EOH concentration was increased an additional 50 mM, and the reaction again plateaued. The equilibrium constant was taken as the average of the six determinations calculated from the two plateaus of the three independently acquired progress curves.

***ROH Redox (Fig S9 B).*** The reaction was monitored *via* increasing RA fluorescence ($\lambda_{ex}$ = 420 nm, $\lambda_{em}$ = 550 nm). The $t_0$ condition: αα-ADH1 (10 nM, active sites), ROH (4.0 μM), $NAD^+$ (10 mM), $KPO_4$ (50 mM), KCl (100 mM), pH 7.4, 25 ± 2 °C. Once the reaction had reached completion, the ROH concentration was increased by an additional 4.0 μM, and the reaction again plateaued. The equilibrium constant was taken as the average of the six determinations calculated from the two plateaus of the three independently acquired progress curves.

##### *GST*

***The DCNB Reaction (Fig S9C).*** Reaction progress was monitored *via* GS-CNB absorbance ($\varepsilon_{345}$ = 9.3 $mM^{-1}$ $cm^{-1}$). Conditions: GSTA1 (5.0 μM, active sites), DCNB (0.30 mM), GSH (0.30 mM), DTT (2.0 mM), and $K_2PO_4$ (50 mM), pH 7.4, 25 ± 2 °C. Once the reaction reached completion, the DCNB concentration was increased by 0.30 mM, and the reaction again plateaued. To assess whether background levels of $Cl^-$ in buffer were sufficiently high to warrant including them in $K_{eq}$ calculations, the $Cl^-$ concentration was determined by monitoring CDNB formation using absorbance ($\varepsilon_{360}$ = 9.3 $mM^{-1}$ $cm^{-1}$)[10]. Conditions: GSTA4 (1.0 μM, active sites), (2,4-dinitrobenzyl)-glutathione (GS-DNB, 5.0 mM)),

DTT (2.0 mM), $KPO_4$ (50 mM), pH 7.4, 25 ± 2 °C. Anion concentrations were determined in triplicate. The $Cl^-$ concentration (6.1 ± 0.5 µM) was included in equilibrium constant calculations described in the preceding paragraph. Fluoride ion contamination, assessed using $^{19}$F-NMR to detect FDNB, was ≤ 500 nM. Approximately one liter of the buffer (DTT (2.0 mM), $K_2PO_4$ (50 mM), pH 7.4) used in the $Cl^-$ concentration determinations was stored in 10 mL aliquots at -80 °C and was used for all GST experiments.

***The 4HNE Reaction (Fig S9D).*** Reaction progress was assessed by monitoring GSH formation. To do so, reactions were quenched by addition of NaOH (100 mM final), boiled (5 min), centrifuged (15,000 x g, 1 min) and GSH in the supernatant was reacted with DTNB to form TNB, which was quantitated using fluorescence ($\lambda_{ex}$ = 412 nm, $\lambda_{em}$ = 550 nm). Reaction conditions: GSTA4 (10 nM, active site), GS-4HNE (5.0 mM), and $K_2PO_4$ (50 mM), pH 7.4, 25 ± 2 °C. Once the reaction reached completion the GS-4HNE concentration was increased by an additional 5.0 mM and the reaction plateaued a second time. The equilibrium constant was taken as the average of the six determinations calculated from the two plateaus of the three independently acquired progress curves.

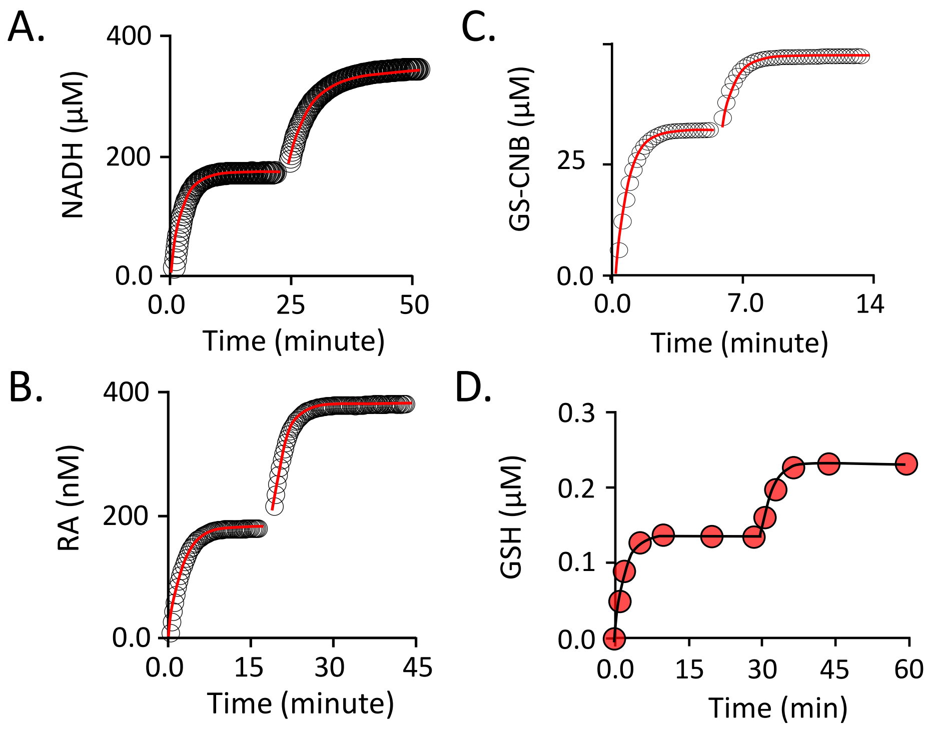

**Fig. S9. Single-Reaction equilibrium constant determinations for ADH and GST catalyzed reactions.** (**A**) *The γγ-ADH1 EOH reaction*. (**B**) *The αα-ADH1 ROH reaction*. (**C**) *The GSTA1 DCNB reaction* (**D**) *The GSTA4 4HNE reaction*. The conditions and protocols associated with Panels A-D are described in the preceding section. In all cases (A-D), Reaction progress curves were least-squares fit to a single exponential equation. The best-fit curves are seen passing through the data and the predicted endpoints were used to calculate equilibrium constants. The second addition of substrate ensures that plateaus are not due to enzyme inhibition or inactivation. All experiments were performed in triplicate. Each data point in Panels A - D is the average of three independent measurements. Each dual-plateau study yields two equilibrium-constant predictions; each triplicate study yields six predictions. The averaged predictions and their errors are compiled in Tables 1 and 3.

#### *GS-4HNE $^{1}H$- NMR*

Linear and cyclic forms of GS-4HNE are distinguishable based on $^{1}$H-NMR of the H1 proton, which transitions from an aldehyde to hemiacetal environment during cyclization. Conditions: GS-4HNE (2.0 mM), $K_2PO_4$ (50 mM), pH 7.4 25 ± 2 °C. $D_2O$ (> 99%) and TSP (0.20 mM) were present in a coaxial NMR sample-tube insert. Spectra were acquired with water suppression using a Bruker 600 MHz spectrometer equipped with a TCI H/F-cryogenic probe[11]. Each spectrum was the average of 1024 scans acquired using a 1D-pulse sequence (20.0 s relaxation delay, 4.0 s acquisition time). The relaxation delay was > than 5.0 times T1 of the cyclic proton (i.e., the proton with the longer T1, 3.2 s). T1 values were determined using a standard T1 *inversion*-recovery protocol[12]. Peaks were fit to a Lorentzian shape using TopSpin3.5[13] and concentrations were determined by normalizing peak areas to that of the TSP signal. Assignment of the cyclic H1 resonance was based on published spectra[14]. The linear H1 resonance was assigned based on its predicted chemical shift and saturation transfer between the linear and cyclic resonances.

***Saturation Transfer Studies***. The forward and reverse rate constants of the GS-4HNE cyclization reaction were determined using $^{1}$H-NMR Transient Saturation-Transfer studies[12] that monitored interconversion of the H1 proton between linear (aldehyde) and cyclic (hemiacetal) forms of GS-4HNE[12,15]. H1 peak intensities of each H1 resonance (linear or cyclic) were measured during on- and off-resonance (-10 ppm) saturation of the partnered resonance over a series of time points. Cyclization rate constants, k, were obtained by least squares fitting to the following equation:

$$I(t) / I_o = [1 / (1 + k \cdot T1)] + [(k \cdot T1) / (1 + k \cdot T1)]\ e^{-t \cdot T1 / (1 + T1 \cdot k)}.$$

I(t) and Io represent the intensity of the non-saturated proton during on- and off-resonance saturation of the partnered proton. Proton relaxation times, T1, were independently determined using T1 inversion-recovery studies (see preceding paragraph). H1 chemical shifts, peak widths at half-height, longitudinal relaxation times (T1), peak suppression at infinite time ($I_{sat}/I_o$) and reaction rate constants (k) are listed in Table S1. Instrumental settings, NMR equipment and solution conditions are described in the preceding paragraph. The saturation data and associated fits are shown in Figs S10 and S1.

**Table S1.** Saturation-Transfer $^1$H-NMR Study GS-4HNE Cyclization

| GS-4HNE State | Chemical Shift (ppm) | Peak Width (ppm) | $T_1$ (s) | $I_{sat}/I_o$ | k ($s^{-1}$) |
|---|---|---|---|---|---|
| Linear | 9.78 | 0.0076 (0.0005) | 1.2 (0.1) | 0.16 (0.03) | 3100 (170) |
| Cyclic | 5.58 | 0.0083 (0.0004) | 3.2 (0.2) | 0.75 (0.01) | 17 (2) |

[a]Parentheses enclose one standard deviation unit.

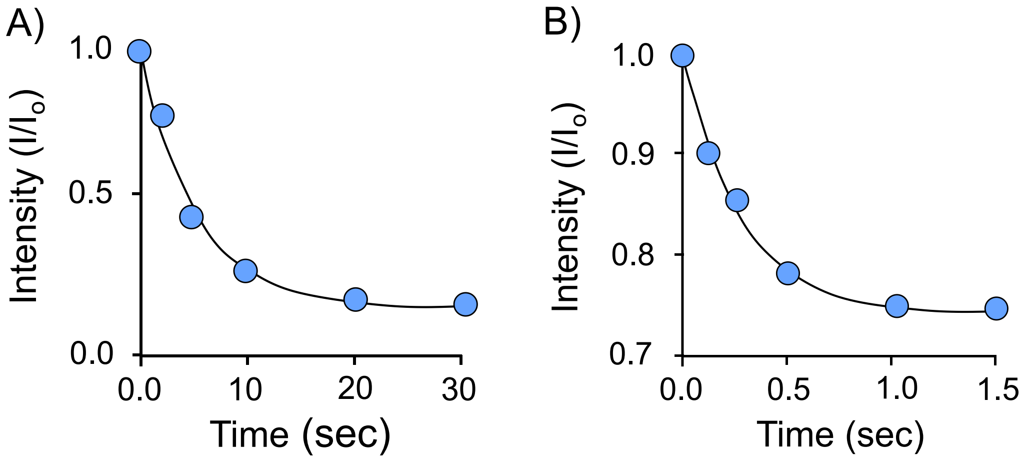


**Fig. S10. Time dependent magnetization transfer between the linear and cyclic forms of the GS-4HNE H1 proton**. (**A**) *Transfer from the cyclic to the linear form.* (**B**) *Transfer from the linear to the cyclic form.* Conditions: GS-4HNE (5.0 mM), $K_2PO_4$ (50 mM), pH 7.4. $D_2O$ (> 99%) and TSP (0.20 mM) were present in a coaxial NMR-tube insert.

##### *The Spontaneous HNE Reaction*

4HNE and GSH slowly react spontaneously to form the conjugate, GS-4HNE[16]. The reaction rate constant is linear with [4HNE][16] and non-linearity with [GSH] has been mentioned[17,18] but not been investigated. To ensure that the non-enzymatic reaction rates were negligible relative to those associate with the initial-rate studies, $k_{obs}$ were determined over a 4HNE x GSH concentration matrix that span the concentrations used in the enzymatic studies and used to compare enzymatic and non-enzymatic rates. The resulting $k_{obs}$-vs-concentration plots, given in Fig S10, reveal that the $k_{obs}$ values, which are consistent with the literature[16-18], are indeed linear with [4HNE] in non-linear with [GSH]. The $k_{obs}$ values were used to calculate the non-enzymatic reaction rates at the [4HNE] and [GSH] used in the initial-rate reactions. Comparison of the rates indicates that the contribution of the spontaneous reaction to the initial-rate measurements ranged from 1.7 – 3.

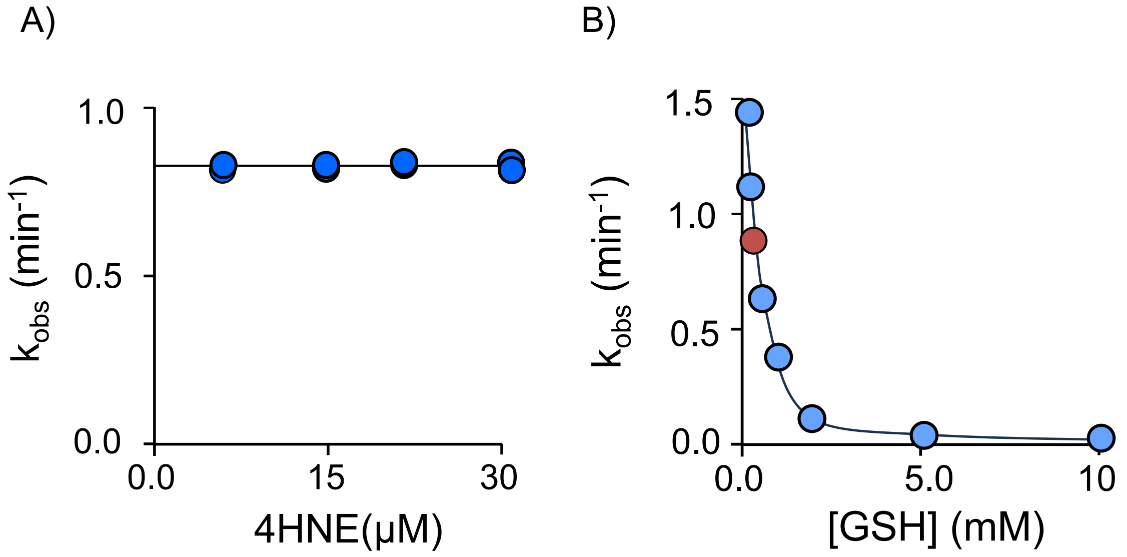


***Fig S11. Spontaneous GSH Conjugation to 4HNE.*** **(A)** *$k_{obs}$ vs [4HNE] at a fixed [HNE]*. **(B)** *$k_{obs}$ vs [HNE] over a series of GSH concentrations*. Reactions were run for > 5 half-lives and rate constants were obtained by least-squares-fitting progress curves to a single exponential. Rate constants were linear with [4HNE] at all [GSH]. Each dot in Panel B represents the average *$k_{obs}$* values obtained from duplicate series of four HNE concentrations — the Panel B red-dot value was calculated from the data seen in Panel A. Conditions: GSH (0.10, 0.20, 0.30, 0.50, 1.0, 2.0, 5.0 and 10 mM), 4HNE (0.020 – 0.10 x [GSH]), $K_2PO_4$ (50 mM), pH 7.4, 25 ± 2 °C. Reactions were monitored *via* the loss of 4HNE absorbance at 225 nm ($\varepsilon_{225}$ = 19500 $M^{-1}$ $cm^{-1}$).

***Modeling.*** Half-site reactions were modeled based on widely accepted mechanisms for the enzymes[1-3]. The models describe energy-coupling, half-site dimers catalyzing two bi-bi substrate reactions[1-3] involving a common reactant. Energy coupling was embedded in the mechanisms by requiring that both subunits turnover before releasing products. GSH and ADH models are presented in Fig S9 and S10. Each step of the mechanism was entered into Gepasi[7], a kinetic simulation program, and assigned a forward and reverse rate constant. All ligand on-rate constants were set at 1 x $10^7$ $M^{-1}$ $s^{-1}$, and off-rate constants were calculated based on $K_m$ values found in Tables 1-4. The on-rate constant was chosen such that all ligand-binding steps are fast enough relative to turnover that all ligand binding steps in the scheme are very near equilibrium. The rate constants governing turnover of central complexes were set equal to experimentally determined $k_{cat}$ values (see, Tables 1-4). The resultant models were used to predict experimental outcomes, seen as solid lines passing through data in Figs 1, 3, and in the selection of experimental conditions.

ADH Mechanism

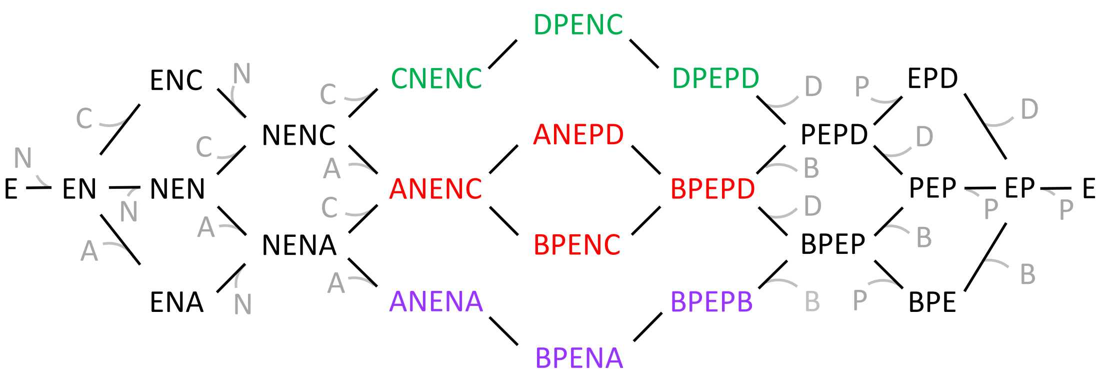


N - $NAD^+$
A - Retinol
C - Ethanol

P - NADH
B - Retinaldehyde
D - Acetaldehyde

**Fig. S12.** ***The ADH Heterodimer Kinetic Scheme***. The scheme is based on the widely accepted ADH sequential-ordered kinetic mechanism. The left and right sides of the symbol ***E*** represent different subunits of the dimer. Green and purple texts indicate complexes unique to homodimer pathways. Red text indicates complexes found only in the heterodimer pathway. Black text indicates complexes common to more than one branch. Grey text indicates solution-phase reactants

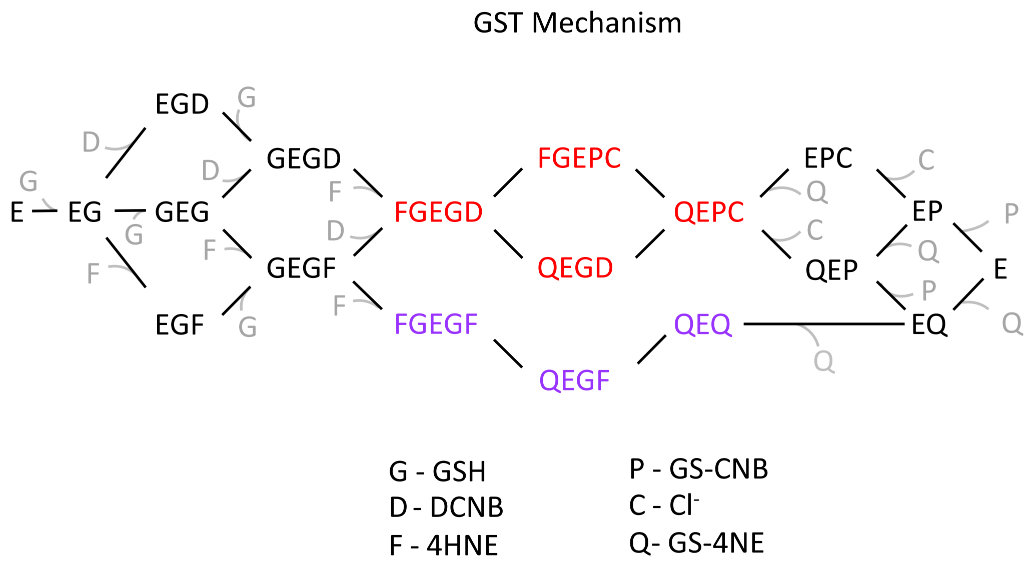


**Fig. S13.** ***Heterodimer GST kinetic scheme.*** The scheme is based on the widely accepted GST sequential-ordered kinetic mechanism. The left and right sides of the symbol *E* represent different subunits of the dimer. Red text indicates complexes found only in the heterocomplex pathway. Purple texts indicate complexes unique to the homocomplex pathway. Black text indicates complexes common to both branches. Grey text indicates solution-phase reactants. There is no DCNB homodimer pathway because DCNB it is not turned over by the heterodimer A4 subunit.

### Supplementary Materials for

***On the Energetics of Small-Molecule Metabolism***

Ian Cook and Thomas S. Leyh*

Half-site enzyme citations were identified in a systematic literature search using web-based tools to query PubMed, PubMed Central, and journal databases (Elsevier, Springer, PLOS One, etc.). The following search terms were used: "half-site reactivity," "half-of-the-sites reactivity," "alternating sites mechanism," "flip-flop catalysis," and "asymmetric dimer catalysis." The top 20 to 30 hits were cross-referenced against review articles and primary literature to verify that each entry met the criteria of experimentally demonstrated half-site or alternating-site behavior, and entries supported only by computational prediction or indirect evidence (e.g., negative cooperativity without burst kinetics or structural asymmetry data) were excluded. Enzymes were categorized by larger families and are presented in the table below. The abstract of each reference was checked by the authors to ensure that half-site reactivity was experimentally demonstrated. While the list is extensive, it should be noted that most enzymes have not yet been tested for half-site reactivity.

**List of Half-Site Reactive Enzymes**

| Metabolic Area / Enzyme Family / Enzyme | Notes | Ref. |
|---|---|---|
| **1. OXIDOREDUCTASES & DEHYDROGENASES** | | |
| Glyceraldehyde-3-phosphate dehydrogenase (GAPDH) | Classic example (S. Bernhard); both positive and negative cooperativity for $NAD^+$ | 1 |
| Human aldehyde dehydrogenase 1A1 (ALDH1A1) | Half-site alternating mechanism | 2 |
| Mitochondrial aldehyde dehydrogenase (ALDH2) | Pre-steady-state burst of 2 mol NADH per tetramer; E487K Asian variant shows dominance | 3 |
| Alcohol dehydrogenase (ADH) | Classic example alongside GAPDH | 4,5 |
| Dihydroorotate dehydrogenase A (DHODA) | Single-molecule fluorescence confirmed half-site signatures | 6 |
| 6-Phosphogluconate dehydrogenase (human erythrocyte) | Binds 2 $NADP^+$ but only 1 NADPH per dimer | 7 |
| Malate dehydrogenase | Shares half-of-sites characteristics | 8 |

| Metabolic Area / Enzyme Family / Enzyme | Notes | Ref. |
|---|---|---|
| Lactate dehydrogenase | Shares half-of-sites characteristics | 8 |
| F420-dependent $NADP^+$ oxidoreductase | First example of half-site reactivity in F420-dependent enzymes; burst kinetics | 9 |
| Glutamate dehydrogenase | Antagonistic homotropic interactions; negative cooperativity for $NAD^+$ | 10,11 |
| Pig lung carbonyl reductase | Half-of-sites reactivity observed | 12 |
| **2. PYRIDINE NUCLEOTIDE DISULFIDE OXIDOREDUCTASES (PNDORs)** | | |
| Glutathione reductase | PNDOR family member with alternating site asymmetry | 13,14 |
| **3. PHASE II DRUG METABOLISM — GLUTATHIONE S-TRANSFERASES (GSTs)** | | |
| Glutathione S-transferase P1-1 (GSTP1-1) | Half-site reactivity protects one Cys47/dimer from nitrosylation; DNDGIC binds one site ($K_i < 10^{-12}$ M) and triggers negative cooperativity at the vacant site ($K_i = 10^{-9}$ M); NO carrier function | 15,16 |
| Glutathione S-transferase A1-1 (GSTA1-1) | Classical half-of-the-sites inhibition by DNDGIC ($K_{i2}$ approaches zero); evolved for DNA protection | 17 |
| Glutathione S-transferase A3-3 (GSTA3-3) | Classical half-of-the-sites inhibition similar to GSTA1-1 | 17 |
| Glutathione S-transferases (Tyr-GST and Ser-GST subfamilies, broadly) | Negative cooperativity with Hill coefficients 0.51–0.75 across 16 tested GSTs; evolved trait present in more recently diverged GSTs but absent in ancestral Cys-GSTs | 17 |
| Human glutathione transferase T2-2 | | 18,19 |
| **4. PHASE II DRUG METABOLISM — SULFOTRANSFERASES (SULTs)** | | |
| Estrogen sulfotransferase (SULT1E1) | First SULT shown to be half-site reactive; burst amplitude = one-half of active sites; product release is rate-limiting | 20 |

| Metabolic Area / Enzyme Family / Enzyme | Notes | Ref. |
|---|---|---|
| SULT2A1 (DHEA sulfotransferase) | Burst of product corresponding to one active-site equivalent per dimer; complete quantitative kinetic model established a catalytic paradigm for the SULT family | 21 |
| SULTs (family-wide) | Three SULTs confirmed to produce pre-steady-state bursts with amplitude of one active-site equivalent per dimer; half-site reactivity is likely paradigmatic for the family | 22 |
| **5. PHASE II DRUG METABOLISM — UDP-GLUCURONOSYLTRANSFERASES (UGTs)** | | |
| UGT2B1 (rat) | Two inactive mutants co-expressed restore activity, proving functional dimerization; amino-terminal domain mediates oligomerization | 23 |
| UGT1A family (broadly) | FRET and co-IP demonstrate homo- and heterodimerization; oligomerization affects enzymatic activity and substrate selectivity. Note: explicit half-site reactivity has not yet been demonstrated as clearly as for SULTs | 24,25 |
| **6. OTHER TRANSFERASES** | | |
| Tyrosyl-tRNA synthetase (*E. coli*) | Classic aminoacyl-tRNA synthetase example | 26 |
| Pig heart L-alanine transaminase | Half-site alternating mechanism | 27 |
| **7. PHOSPHATASES & KINASES** | | |
| Intestinal alkaline phosphatase | One of the earliest characterized half-site enzymes; flip-flop mechanism model (Lazdunski) | 28 |
| Fructose-1,6-bisphosphatase | Demonstrated by quench-flow and isotope trapping; $Mg^{2+}$-dependent | 29 |
| **8. NUCLEOTIDE METABOLISM ENZYMES** | | |
| Thymidylate synthase (TS) | Structural mechanism via relay of conformational changes between | 30,31,32 |

| Metabolic Area / Enzyme Family / Enzyme | Notes | Ref. |
|---|---|---|
| | subunits | |
| Thymidylate synthase–dihydrofolate reductase (TS-DHFR, bifunctional) | Parasitic bifunctional enzyme | 31,32,33 |
| Cytidine triphosphate synthetase (CTPS) | Affinity label binds half of subunits but abolishes all glutamine activity | 34,35 |
| Ribonucleotide reductase (Class Ia, *E. coli*) | Asymmetric β subunit binding revealed by cryo-EM; radical transfer pathway active in only one site | 36 |
| Purine nucleoside phosphorylase (PNP, calf spleen) | "Third-the-sites" reactivity variant; tight-binding hypoxanthine intermediate | 37 |
| DAHP synthase (3-deoxy-D-arabino-heptulosonate 7-phosphate synthase) | Half-of-sites inhibition; inter-subunit communication | 38 |
| **9. CARBOXYLASES & DECARBOXYLASES** | | |
| Biotin carboxylase (bacterial) | Substrate-induced synergism and half-site reactivity | 39 |
| Pyruvate carboxylase | Half-site alternating mechanism for biotin translocation between BC and CT domains | 40 |
| Human pyruvate dehydrogenase | | 41,42 |
| α-Amino-β-carboxymuconate-ε-semialdehyde decarboxylase (ACMSD) | Zinc-dependent; observable differences in metal-ligand positioning between paired sites; Arg239 contributed from adjacent subunit | 8 |
| **10. OXYGENASES & DIOXYGENASES** | | |
| *p*-Hydroxyphenylpyruvate dioxygenase (HPPD) | | 43 |
| Tryptophan 2,3-dioxygenase (TDO) | Heme-dependent; kynurenine pathway | 8 |
| Prostaglandin endoperoxide H synthase 1 (PGHS-1 / COX-1) | Conformational heterodimer throughout catalysis | 44 |
| Prostaglandin endoperoxide H synthase 2 (PGHS-2 / COX-2) | Heterodimer studies proved half-of-sites; pharmacological target of NSAIDs | 44 |
| **11. AMINE OXIDASES** | | |
| Bovine serum amine oxidase | Flip-flop catalysis reported in | 45,46 |

| Metabolic Area / Enzyme Family / Enzyme | Notes | Ref. |
|---|---|---|
| | biochemical studies | |
| **12. ISOMERASES** | | |
| Heptose isomerase (GmhA) | Achieves half-site reactivity without conformational changes; water channel couples paired sites | 47 |
| Glyoxalase I | Zinc-dependent; NMR shows inequivalent active sites | 8 |
| **13. LYASES & SYNTHASES** | | |
| Malate thiokinase | Half-site alternating mechanism | 48 |
| **14. DESATURASES** | | |
| Castor Δ9-18:0-acyl carrier protein (ACP) desaturase | Heterodimers with one active site show full wild-type activity | 49 |
| **15. AMINO ACID BIOSYNTHESIS ENZYMES** | | |
| Aspartate β-semialdehyde dehydrogenase (ASADH) | Extensive interchain contacts create half-site behavior; antimicrobial drug target | 50 |
| Glutamine:fructose-6-phosphate amidotransferase (GFAT/GlmS) | Half-site occupancy observed crystallographically | 51 |
| **16. AMINOMUTASES** | | |
| Lysine 2,3-aminomutase | Kinetic and spectroscopic evidence for negative cooperativity and half-site reactivity | 52 |
| **17. ADENYLYLTRANSFERASES** | | |
| Nicotinate mononucleotide adenylyltransferase (NMAT, *B. anthracis*) | X-ray structural evidence for negative cooperativity in substrate binding | 53 |
| **18. E3 UBIQUITIN LIGASES** | | |
| CHIP E3 ubiquitin ligase | Symmetry breaking during homodimeric assembly activates the ligase; functional asymmetry | 54 |
| **19. VIRAL PROTEASES** | | |
| SARS-CoV-2 main protease (Mpro/3CLpro) | Single-ligand occupancy and negative cooperativity observed; unexpected half-site behavior | 55 |
| **20. ELECTRON TRANSPORT CHAIN** | | |

| Metabolic Area / Enzyme Family / Enzyme | Notes | Ref. |
|---|---|---|
| Dimeric cytochrome *bc*$_1$ complex | Half-of-sites for ubiquinol oxidation; regulated by inhibitor binding at ubiquinone reduction site | 56,57 |
| **21. REDUCTIVE CARBOXYLASES** | | |
| Crotonyl-CoA carboxylase/reductase (CCR) | Intersubunit coupling enables fast $CO_2$ fixation; half-site behavior observed via XFEL crystallography | 58 |
| **22. TPP-DEPENDENT ENZYMES (FLIP-FLOP MECHANISM)** | | |
| Transketolase | Dynamic nonequivalence of active sites; flip-flop mechanism common to TPP-dependent enzymes | 59 |
| Pyruvate decarboxylase | Flip-flop catalysis reported in biochemical studies | 60 |
| **23. DEHALOGENASES** | | |
| Fluoroacetate dehalogenase (FAcD, *Rhodopseudomonas palustris*) | Even under gross excess of substrate, turns over only one substrate at a time; apo crystal structure shows pre-existing asymmetry; empty protomer contributes via enhanced dynamics and water egress | 61 |
| **24. MERCURY REDUCTASES** | | |
| Mercuric ion reductase (MerA) | Alternating sites model; pyridine nucleotide-complexed dimers are asymmetric; | 62 |
| **25. DECAPPING ENZYMES** | | |
| DcpS scavenger decapping enzyme | Crystal structures show symmetric apo-form but asymmetric ligand-bound state; one active site closed with ligand, the other open | 63 |
| **26. FATTY ACID SYNTHASES** | | |
| Animal fatty acid synthase (FAS) | Engineered heterodimer with one wild-type and one fully inactivated subunit is full inactive; | 64 |
| **27. PROTEINS WITH HALF-SITE LIGAND BINDING (NON-ENZYMATIC)** | | |
| Insulin hexamer | Displays half-site reactivity, negative cooperativity, and positive cooperativity for phenolic ligands; quantitative SMB model | 65 |
| Aspartate chemoreceptor (*E. coli* and *S.* | Binds ligand with negative and half- | 66 |

| Metabolic Area / Enzyme Family / Enzyme | Notes | Ref. |
| --- | --- | --- |
| *typhimurium*) | of-the-sites cooperativity | |
